# Diel thermal variability does little to mitigate bleaching in two endangered coral holobionts

**DOI:** 10.64898/2026.09.11.750923

**Authors:** Shelby E. Gantt, Nicola G. Kriefall, Hannah E. Aichelman, Alexa K. Huzar, Clara E. DiVincenzo, Emily Madish, Hanny E. Rivera, Sarah W. Davies

## Abstract

Research has shown that tropical corals inhabiting warmer, more thermally variable reef environments often exhibit reduced bleaching. However, the influence of diel thermal variation (DTV) is rarely tested experimentally independent of changes in mean temperature, and few studies have explored responses of coral hosts together with their associated microbial consortia (Symbiodiniaceae algae and bacteria). Here, using a common garden approach with two species of endangered reef-building corals (*Orbicella franksi* & *O. faveolata*), we show that 90 days of DTV priming (26°C ± 2.4°C day^-1^) elicited little physiological or gene expression plasticity and did not alter Symbiodiniaceae or bacterial compositions relative to corals under stable conditions (26°C). Critically, when these corals experienced a subsequent heat challenge (32°C for 15 days), both species exhibited broad physiological and gene expression responses regardless of DTV priming. A systems biology approach linking gene expression with experimental treatments and traits supports the presence of both 1) convergent gene modules associated with homeostasis in both host species and 2) divergent gene modules showcasing enrichment of catabolism in *O. faveolata* and classic protein folding functions in *O. franksi* under thermal challenge. Together, these data highlight that while congeneric coral species exhibit divergent gene expression responses to thermal challenges, DTV priming did not facilitate thermal stress resistance. These findings suggest that other environmental parameters that co-vary with DTV on a reef may play a more influential role in bleaching mitigation.

## INTRODUCTION

Global temperatures are rising due to environmental change, and seasonal thermal variability is also expected to increase, contributing to more frequent and extreme thermal events (Bathiany et al., 2018; Destri et al., 2025; Thornton et al., 2014). These thermal extremes push organisms to adapt and acclimate at a faster pace than in previous centuries (Kroeker et al., 2020; Thornton et al., 2014), especially in marine environments (Pinsky et al., 2019). Most experimental studies of thermal acclimation in marine organisms have manipulated mean temperatures, yielding valuable insights into plasticity and physiological limits. However, recent work has revealed the importance of diel thermal variability (DTV) in driving organismal responses to environmental change (Klein et al., 2019; Kroeker et al., 2020; Rivest et al., 2017). Therefore, incorporating realistic DTV into experimental designs is critical to improve predictions of species’ responses to climate change (Dee et al., 2020; Sheldon & Dillon, 2016).

Reef-building corals are particularly vulnerable to temperature changes, as ocean warming causes coral bleaching (the breakdown of coral-algal symbiosis) and associated mortality (Brown, 1997; Lesser, 2011; Reimer et al., 2024). In response, research has focused on identifying factors that confer thermal tolerance. Corals from warmer, thermally variable environments often exhibit enhanced heat tolerance compared to conspecifics from cooler, more stable reefs (Kenkel & Matz, 2016; Palumbi et al., 2014; Safaie et al., 2018), though there are exceptions (*e.g.* Klepac & Barshis, 2020). Because temperature covaries with factors such as nutrients, light, and dissolved oxygen across naturally variable reef habitats (Cyronak et al., 2020; Pezner et al., 2021), isolating the role of DTV is challenging. While some *ex situ* studies have manipulated DTV without also altering mean temperature (Aichelman et al., 2025; Jiang et al., 2017; Schoepf et al., 2022), many direct and mechanistic effects of DTV on corals remain unresolved.

Corals are complex holobionts, with many tropical species forming obligate symbioses with Symbiodiniaceae algae and associations with bacterial, archeal, and viral partners (Knowlton & Rohwer, 2003; LaJeunesse et al., 2018; Thompson et al., 2014). Symbiodiniaceae have been established as key players in resistance and resilience to coral bleaching, with dominant types often varying markedly by local environmental conditions (Gantt et al., 2024; Howells et al., 2020; van Oppen et al., 2018) and in response to thermal anomalies (Starko et al., 2023; Van Nynatten et al., 2025). Associations with the Symbiodiniaceae genus *Durusdinium* have also been observed in more environmentally variable habitats (Gantt et al., 2024; Oliver & Palumbi, 2011) and have been associated with host heat tolerance (Manzello et al., 2019; Stat & Gates, 2011) (but see Abrego et al., 2008; Fifer et al., 2026). While the role of the bacterial microbiome is comparatively less established, recent work indicates bacterial partners can also influence host fitness (Doering et al., 2021; Rosado et al., 2019; Santoro et al., 2021) and partner identities are shaped by local environments (Gantt et al., 2024; Kriefall et al., 2022; Ziegler et al., 2017). However, there are also examples of negative interactions with Symbiodiniaceae and bacteria that lead to loss of fitness in coral hosts (D. M. Baker et al., 2018; Vega Thurber et al., 2020), highlighting the complexity of interactions underlying phenotypes.

Coral hosts within the genus *Orbicella* offer a well-studied system to explore holobiont responses to environmental variation. *Orbicella* spp. are ecologically critical reef-builders in the Caribbean that have experienced substantial declines and are recognized as threatened (NOAA, 2023). Considered life-history “generalists” that perform well across a range of environments (Darling et al., 2012), members of the *Orbicella* species complex can differ in corallite spacing, physical structures, and niche partitioning (Knowlton & Jackson, 1994), and previous work showed species-specific responses to reciprocal transplantation (Prada et al., 2022). *Orbicella* spp. also associate with diverse microbiota, including multiple Symbiodiniaceae genera whose relative abundances can shift under heat stress (Cunning et al., 2015; Kemp et al., 2014, 2015). Their bacterial communities are also comparatively well-characterized (Prada et al., 2022; Sunagawa et al., 2009) and their gene expression responses to various stressors have been extensively explored (Anderson et al., 2016; Pinzón et al., 2015; Schlecker et al., 2022; Strader et al., 2024; Wright et al., 2019). Collectively, species-specific differences within this species complex provide a framework to explore how dynamic thermal regimes influence corals of urgent conservation concern.

To assess whether DTV can influence plasticity in holobiont traits and enhance coral thermal tolerance, we exposed two *Orbicella* species to fluctuating versus stable thermal priming conditions, followed by a heat challenge. Specifically, clonal fragments of *Orbicella faveolata* and *O. franski* were maintained in aquaria under either fluctuating (26°C ± 2.4°C day^-1^) or stable (26°C) regimes for 90 days of thermal priming, after which we profiled gene expression, Symbiodinaceae and bacterial communities, and various physiological metrics. Next, fragments were subjected to a heat challenge (32°C) for 15 days and the same holobiont traits were re-profiled. DTV priming produced minimal shifts in symbiotic associations and holobiont physiology. Some species-specific host responses and shifts in Symbiodiniaeae gene expression were observed during the heat challenge, but DTV priming did not alter holobiont heat stress responses in either coral species. Although field-based studies have linked DTV to bleaching mitigation, we conclude that other co-varying environmental parameters across reefs may better explain this phenomenon.

## METHODS

### Coral collection

In August 2018, large fragments (∼20 cm^2^) of *Orbicella faveolata* (*n* = 5) and *O. franksi* (*n* = 5) colonies were collected *via* scuba diving at ∼23 m depth from the east bank of the Flower Garden Banks (FGB) National Marine Sanctuary (27°57’31.68”, 93°38’33.81”; permit FGBNMS-2018- 006). Colonies were transported to Boston University, fragmented with a band saw into ∼4 cm^2^ pieces, and glued to ceramic tiles. Tanks (400 L, ∼26°C, and 33-35 ppt salinity) were maintained for ∼2 years with weekly water changes (∼5–20%) and water quality monitoring (nitrates, phosphates, calcium, magnesium, and alkalinity). Light levels (AquaIllumination Hydra) were maintained at ∼40 µmol*m^−2^s^−1^ with a 10:14 h light:dark cycle, where light hours were flanked by one-hour ramps to simulate sunrise and sunset. Corals were fed twice weekly with Reef-Roids (PolypLab) and once weekly with live brine shrimp (*Artemia*).

### Experimental design & tank conditions

In July 2020, six fragments randomly selected from each colony were halved, yielding 12 fragments per colony. Subsequent genotyping revealed that the ten colonies represented only four unique genets per species, as one pair of clones was identified within each species (Figure S1; see sequencing methods below). Fragments were randomly assigned to one of six 75 L tanks, of which three were maintained with stable conditions and three with diel thermal variability (DTV), for a total of six replicate fragments per genet per treatment (Figure 1). Corals recovered from fragmentation at conditions described above for 11 days. Water changes (∼20%), water quality assessments (described above), algae scrubbing, and fragment rotations (to reduce microhabitat differences) were performed weekly. Salinity measurements were taken five days per week. Pendant HOBO loggers recorded temperature every 15 min, with one logger deployed per treatment.

**Figure 1.**
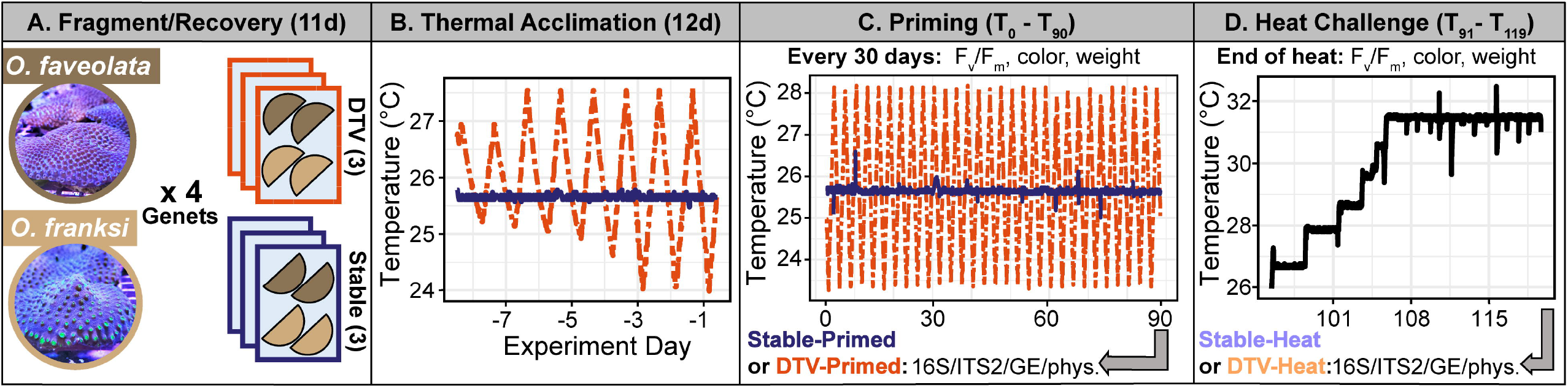
Experimental design overview. Diagram representing (A) fragmentation and placement of *Orbicella faveolata* (brown) and *O. franksi* (beige) into stable (blue) or DTV (orange) thermal priming treatments. (B) Thermal acclimation of fragments to DTV or stable thermal priming treatments. (C) Thermal regimes and sampling for thermal priming experiment. One Stable-Primed (blue) and one DTV-Primed (orange) fragment per genet was sampled on experimental day 90. (D) Heat challenge experiment showing sample timing of one Stable-Heat (light blue) and one DTV-Heat (light orange) fragment per genet. Grey arrows in C and D indicate when samples were frozen for downstream analyses (T_90_ and T_119_).

After recovery, DTV tanks were ramped from 26°C ± 0°C day^-1^ to 26°C ± 2.4°C day^-1^, with temperatures increasing and decreasing by 0.6°C every three days for 12 days using the Apex Aquacontroller (Neptune Systems; Figure 1). Initial ramping was uneven due to a chiller issue; however, this did not continue into the thermal priming period, which was maintained for 90 days. Both temperature regimes were based on long-term monitoring data collected at the west FGB using subsurface temperature recorders at 25 m depth from 2015–2019 (Enochs et al., 2024). The stable treatment was maintained at 26°C to correspond to the annual mean temperature from this period, while the DTV treatment was centered on the same mean and programmed to a daily range of 4.8°C, which was similar to the maximum observed diel range (∼4.4°C) within technical constraints. Corals in both treatments began to pale by day 34, so light levels were reduced to ∼20 µmol*m^−2^s^−1^ on an 8:16 h light:dark cycle for the remainder of the study. Since lights for all tanks were shifted in parallel, these changes do not affect comparisons between priming treatments. After 90 days, 30 fragments per treatment (*n* = 60 total) were flash frozen in liquid nitrogen and maintained at -80°C for downstream processing. For the heat challenge, the remaining fragments (*n* = 60) were consolidated into three tanks, ramped from 26°C to 32°C over 10 days, and held at 32°C for 15 days (Figure 1). Fifteen days simulates an extended heatwave and 32°C is a common heat challenge threshold in *O. faveolata* (Cunning et al., 2015; Grottoli et al., 2014). At T_119_, all remaining fragments were flash frozen and maintained at -80°C.

Unless otherwise stated, all analyses used R version 4.4.1 (R Core Team, 2024), and scripts are available at https://github.com/shegantt/DTV_Ofav_Ofra. Mean temperature and concentrations of nitrate, phosphate, calcium, alkalinity, and magnesium were compared using a Wilcoxon rank sum test, as data failed to meet parametric test assumptions, while salinity was compared *via* a one-way ANOVA. For all water quality metrics, Levene’s tests checked for homogeneity of variance between treatments.

### Coral holobiont physiological traits

The following non-invasive physiological metrics were collected during thermal priming every 30 days (T_0_, T_30_, T_60_, T_90_) for the stable- or DTV-primed fragments and after the heat challenge (T_119_; “Stable-Heat” or “DTV-Heat” fragments): photosynthetic efficiency (F_v_/F_m_) measured using a junior Pulse-Amplitude-Modulation chlorophyll fluorometer (Walz), fragment images using an iPhone SE, and buoyant weight (Spencer Davies, 1989). To estimate coral color, images were white-balanced in Adobe Photoshop and red channel intensity was quantified in MATLAB by averaging ten random points across the coral surface, a common proxy for Symbiodiniaceae pigment density (Winters et al., 2009). Images from the end of thermal priming and heat challenge were also used to estimate fragment surface area in triplicate with ImageJ (Schneider et al., 2012). For T_90_ and T_119_ fragments, small (∼0.5 cm^2^) tissue samples were excised, preserved in 200 proof ethanol, and stored at -80°C until sequencing preparation for DNA and RNA analyses.

For physiological analyses, replicate F_v_/F_m_, buoyant weight, and red channel intensity measurements were averaged for each fragment and genotype. Buoyant weights were calculated as the difference between consecutive time points (*i.e.*, T_30_-T_0_, T_60_-T_30_, etc.) transformed to percent change, hereafter termed “growth rate”. To ensure that weight changes were being compared across similar timescales, T_90_-T_60_ was used to test differences between thermal priming treatments and T_119_-T_90_ was used to test how heat challenge influenced growth. For each species, F_v_/F_m_, red channel intensity, and growth rate were compared between thermal priming treatments and time points using a linear mixed effects regression (lmer) model (Metric∼Priming*Time point) with a random effect of genotype using lme4 (Bates et al., 2014). The *check_model* function in performance (Lüdecke et al., 2021) tested model assumptions. This analysis was carried out twice: once testing differences between T_0_ and T_90_ to determine the effect of thermal priming, and the second analysis using only fragments sampled after thermal priming (T_90_) and heat challenge (T_119_) time points. Pairwise comparisons between interaction terms (Priming:Time point) were then tested using emmeans (Lenth et al., 2022) with Bonferroni-correction.

Photosynthesis (gross and net) and dark-adapted respiration rates were measured on day 91 by quantifying dissolved oxygen (O_2_) concentrations in 624 mL acrylic chambers using a 10- channel Fiber Optic Oxygen Transmitter with dipping probes and integrated temperature probe (OXY-10 SMA, Pre-Sens Precision Sensing GmbH) and Pre-Sens Measurement Studio 2 software (version Oxy10v3_33fb) at 0.07 Hz, which automatically corrected for temperature and salinity. A ten-chamber system with built-in stir plates (Australian Institute of Marine Science) submerged in a recirculating water bath of seawater (∼33 ppt) was maintained at 26°C, with one coral fragment in each of nine chambers. A blank chamber (seawater only) was included in all runs to account for background microbial metabolism and instrument drift. O_2_ evolution was recorded every 15 seconds for 20 minutes in the dark to obtain dark-adapted respiration rates (R) and then under light (AI Hydra Twentysix HD; ∼20 µmol*m^−2^s^−1^) for 20 minutes to quantify gross photosynthesis (P). This procedure was repeated after the heat challenge (T_118_), with identical parameters except the water bath was maintained at 32°C. R and P rates were calculated from O_2_ concentrations using the LoLinR package with smoothing parameter of 0.4 (Olito et al., 2017). Rates were blank corrected and normalized to chamber volume and fragment surface area. Differences in P, R, and P/R ratios across treatments were completed as above for physiology.

### Symbiodiniaceae & bacterial community characterization

To characterize Symbiodiniaceae and bacterial communities, one fragment from each genet (*n* = 5/species, prior to clone identification) and thermal priming treatment was selected at random after 90 days (*n* = 20) and heat challenge (*n* = 20). Total DNA was extracted using the RNAqueous™ Total RNA Isolation Kit (Thermo Fisher) before DNAse I application. Primers *SYM_VAR_5.8S2* and *SYM_VAR_REV* (Hume et al., 2018), with modifications as in Baumann et al. (2018), were used to amplify the Symbiodiniaceae-specific ITS2 region as in Aichelman et al. (2025). Briefly, PCR mix included 0.1 µL DNA template, 0.025 U ExTaq with 1x ExTaq buffer (Takara), 1 µM primers, 0.2 mM dNTPs, and molecular-grade water up to a volume of 20 µL. Cycling conditions were 95°C for 5 min; 30 cycles of 95°C for 1 min, 59°C for 2 min, and 72°C for 2 min; and a final extension of 72°C for 10 min. Amplicons were cleaned with a GeneJet PCR Purification Kit (Thermo Fisher), dual-indexed in a second PCR (6 cycles with the same conditions), and pooled by relative band intensity on a 1% agarose gel. The final library was cleaned again, excised from a 2% agarose gel, and quantified on a spectrophotometer (DeNovix).

16S amplicon library preparation followed ITS2 specifications, except primers *Hyb515F* (Parada et al., 2016) and *Hyb806R* (Apprill et al., 2015) amplified the V4 region of the 16S rRNA gene with a 62°C annealing temperature. One *O. faveolata* sample failed to amplify (Table S2). 16S libraries, including two negative controls, were pooled with ITS2 libraries at a ratio of 1:2 ITS2:16S. Sequencing (250 bp, paired-end) was conducted on an Illumina MiSeq at Tufts University Core Facility.

Bbmap (Bushnell, 2014) was used to retain reads beginning with ITS2 primer sequences and these data were submitted to SymPortal (Hume et al., 2019) to identify ITS2 type profiles. Average raw reads per sample were 36,667 (± 11,103) and 23,465 (± 8,015) post-quality control by Symportal (Table S2). After clone identification (see below), one representative from each clone pair was removed at random. Counts assigned to type profiles were transformed to relative abundances in phyloseq (McMurdie & Holmes, 2013), and type profile counts were summed by the ITS2 sequence that made up the majority of each profile (“Majority_ITS2_Sequence”). ITS2 type “E1c” was present in only one sample with < 0.01% relative abundance, thus it was removed from further analysis.

To test whether ITS2 types differed by experimental variables, a generalized linear mixed model (GLMM) was fitted using the glmmTMB package (Brooks et al., 2017). As the same type profiles appeared in both coral species, and to increase statistical power, data from both species were combined into one model. The full model included fixed effects of host species and the interaction between thermal priming and heat challenge, as well as a random effect of host genotype and ITS2 type identity, assuming an ordered beta distribution (logit link function). Additional random effects were included to allow ITS2 type identity to interact with all aforementioned variables. The full model was compared to reduced models lacking interactions between ITS2 type and thermal priming with heat challenge, or between ITS2 type and host species, using log-likelihood ratio tests assuming a ^2^ distribution, to test whether ITS2 proportions changed according to these variables.

Bbmap (Bushnell, 2014) retained raw reads containing the 16S primer and cutadapt (Martin, 2011) removed primer sequences. Average raw reads per sample were 50,073 (± 30,016; Table S2). DADA2 (Callahan et al., 2016) inferred amplicon sequence variants (ASVs), and ASVs whose lengths were outside of 244–264 bp were removed. Taxonomy was assigned using the Silva v138 database (Quast et al., 2013) and ASVs identified as chloroplasts (*n* = 237), mitochondria (*n* = 22), or non-bacterial kingdoms (*n* = 31) were removed. We used decontam (Davis et al., 2018) and the two negative controls to remove five ASVs identified as putative contaminants. ASVs present in only one sample and those making up less than 49 reads total (*i.e.* < 0.01% of the sequencing library’s total read counts) were removed. Counts less than 10 within a sample were zeroed (see Nikodemova et al., 2023), leaving a final total of 465 ASVs and an average of 12,689 (± 10,464) reads per sample. After clone identification (see below), only one representative from each genotype was retained for further analysis (Table S2).

To compare alpha diversity across treatments, phyloseq (McMurdie & Holmes, 2013) calculated Shannon and Simpson’s indices. For each host species, these diversity metrics were compared across thermal priming treatments and heat challenge with a linear mixed effects regression (Metric∼Priming*Heat), including a random effect of genotype, using the lme4 package (Bates et al., 2015). *Post hoc* pairwise comparisons were conducted using emmeans (Lenth et al., 2022). Diversity metrics were re-analyzed following rarefaction down to 3,149 reads to ensure results were not an artifact of uneven sampling depth. Sample relative abundance data (generated from the non-rarefied dataset) were then used to compare Bray-Curtis dissimilarity between samples. Communities across treatment groups were compared using the same approach as for Symbiodiniaceae, except that the observed counts (rather than proportional data) were analyzed; accordingly, models assumed a negative binomial type I error distribution (log link function) and included a term that offset the model by the logarithm of the total read counts for each sample to account for uneven read depth.

### Host and Symbiodiniaceae gene expression analyses

Total holobiont genomic material extracted from thermally primed (*n* = 20) and post-heat challenge (*n* = 20) samples described above were treated with DNase I (Invitrogen), normalized, and submitted to the Genomic Sequencing and Analysis Facility at UT Austin for TagSeq (Meyer et al., 2011) standard coverage library preparation and NovaSeq S1 SR100 sequencing.

A total of 282.4 million reads were generated from 40 samples, with an average of ∼7.1 million reads per sample (5.2-9.2 million reads/sample; Table S2). Fast-X toolkit trimmed Illumina adaptors and removed poly-A tails, sequences < 20 bp in length, and sequences with Phred quality scores < 20. PCR duplicates were removed, leaving 73.4 million reads (0.8-2.6 million reads/sample). To confirm sample identification and identify clones, single nucleotide polymorphisms (SNPs) were identified by mapping quality-filtered reads to the *Orbicella faveolata* genome (Prada et al., 2016) using Bowtie v2.2.0 (Langmead & Salzberg, 2012). 48,194 SNPs were identified and ANGSD (Korneliussen et al., 2014) calculated pairwise identity-by-state (IBS) matrices using a minIndDepth filter of 5. Hclust visualized individual relatedness, which identified two sets of clones, one per species (Figure S1). One of each clone pair was removed from analysis (VB for *O. faveolata*, KA for *O. franksi*).

To confirm ITS2-based Symbiodiniaceae types, filtered reads were mapped using Bowtie2 v2.5.1 (Langmead & Salzberg, 2012) to a holobiont reference, including the *O. faveolata* genome (Prada et al., 2016), and a database of Symbiodiniaceae transcriptomes (*Symbiodinium* sp. and *Breviolum* sp. from Bayer et al. (2012); *Cladocopium* sp. and *Durusdinium* sp. from Ladner et al. (2012)), following the pipeline in Barfield et al. (2018). Reads mapping to the Symbiodiniaceae database were used to calculate relative abundances of each genus. Both ITS2 and TagSeq SNP analyses showed that all corals hosted *Durusdinium*, with ITS2 data indicating 15/16 *O. faveolata* and 12/16 *O. franksi* fragments were dominated (> 50% relative abundance) by *Durusdinium*, while SNP analyses indicated all 16 *O. faveolata* and 13/16 *O. franksi* fragments were dominated by *Durusdinium* (Figures 3B, S2).

Due to the relative *Durusdinium* dominance, filtered holobiont reads were remapped to a holobiont genome consisting of concatenated *O. faveolata* (Young et al., 2024) and *D. trenchii* (Dougan et al., 2024) genomes. Resulting count files were split into host- and symbiont-specific files. Host counts yielded 0.45–1.50 million reads per sample (mean ± SE: 1,020,437 ± 37,890; Table S2). Host counts were filtered to exclude genes with mean counts of < 2 in 90% of samples, resulting in 16,447 host genes. DESeq2 v1.44.0 (Love et al., 2014) identified differentially expressed genes (DEGs) using the design ∼HostSpecies+Priming*Heat with an FDR-correction of *p* < 0.10.

Next, using the DESeq model design ∼1, combined host counts were *rlog-*normalized and a principal component analysis (PCA) was conducted using *prcomp* to characterize differences in gene expression (GE) between thermal priming and heat challenge, which were tested using a PERMANOVA with the *adonis2* function in vegan v2.6-10 (Oksanen et al., 2013). Next, significance in host species-specific GE across treatments was tested and pairwise differences were determined using the *pairwise.adonis2* function (Martinez Arbizu, 2020). Pairwise differences were tested between overall GE for both host species combined and for each host species separately. DESeq2 identified DEGs using the design ∼HostSpecies+TreatHeat with TreatHeat as a factor combining thermal priming and heat challenge into one factor to allow pairwise comparisons across all groups. Shared host DEGs by thermal priming and heat challenge were identified using the ggvenn package (Yan, 2025).

All host genes were used as input for a weighted gene co-expression network analysis (WGCNA; Langfelder & Horvath, 2008) to identify gene modules associated with treatments and traits. Briefly, signed gene co-expression networks were constructed for host species combined and separately (parameters: softpower = 5; minimal module size = 35; deepsplit = 0; MEdiss = 0.45). Module eigengenes were correlated with holobiont traits (F_v_/F_m_, red channel intensity, buoyant weights, P, R, and P/R) and treatments (DTV-Primed, Stable-Primed, DTV- Heat, Stable-Heat). Modules correlating with treatments and traits for the combined analysis (*n* = 15) and individual host species (*O. faveolata* = 14, *O. franksi* = 14) were visualized in a module-trait heatmap.

Gene ontology (GO) enrichment analysis was performed on modules showcasing significant correlations with traits or treatments. Each gene within a module received a module membership (kME) value, representing the strength of its association and genes not in that module were given zeros. GO enrichment analyses were performed for each GO category (biological processes (BP), cellular function (CC), and molecular function (MF)) using Mann-Whitney U tests (Wright et al., 2015). We quantitatively compared the two ‘brown’ modules identified within both host species showing similar GO enrichment with a semantic similarity matrix using GOSemSim (Yu et al., 2010). The resulting similarity matrix was visualized with a heatmap using Euclidean distances and complete linkage. Next ‘brown’ modules were filtered for shared GO terms, adjusted p-values were extracted, –log10(padj) transformed, and fitted to a linear model to estimate slope and R² using a Kendall rank correlation. Shared significantly enriched MF GO terms between the two ‘brown’ modules were visualized using the ggvenn package (Yan, 2025).

Symbiodiniaceae counts yielded 84,708–1,169,007 reads (mean ± SE: 510,693 ± 48,919) per sample (Table S2). Symbiodiniaceae-specific analyses included DESeq2, PCA, and PERMANOVA analyses performed as described above for host analyses. After filtering, 2,188 genes remained, which was an insufficient number of genes to conduct WGCNA; however, PCA identified significant overall expression differences across treatment groups. To explore whether GE plasticity differed across different treatments, the *PCAplast* function (Bove et al., 2022) calculated weighted distances that each fragment moved in each treatment (DTV-Primed, DTV- Heat, Stable-Heat) relative to its Stable-Primed (T_90_) control. An ANOVA followed by Tukey’s HSD tests compared groups after assumptions were tested with Shapiro-Wilk’s tests and the *check_model* function. To test for differences in dispersion of GE plasticity among samples within groups, the parametric Levene’s test (L) and non-parametric Filgner’s test (F) were used. Coral host species was not included as a factor in any symbiont models because this factor was a significant driver of Symbiodiniaceae GE in the PCA. GO enrichment analyses were conducted using –log10(padj) values from DEGs across pairwise treatment contrasts and overall thermal priming (Stable vs. DTV) and heat challenge (Primed vs. Heat) comparisons; however, no significant GO terms were identified for any comparison.

## RESULTS

### Experimental water quality during thermal priming

During priming, mean temperature (± SD) was significantly higher in DTV-primed tanks (25.73 ± 1.36°C) than stable-primed tanks (25.67 ± 0.16°C; *p* < 0.001); however, the effect size was small (r = 0.03, Table S1; HOBO logger accuracy ± 0.2°C). Mean daily range during priming was significantly higher (4.74 ± 0.02°C) in DTV-primed tanks compared to stable-primed tanks (0.33 ± 0.03°C; *p* < 0.001; Table S1). Means and variance of salinity, nitrate, phosphate, magnesium, calcium, and alkalinity did not differ between priming treatments or heat challenge (Table S1).

### DTV priming and heat challenge influence holobiont physiology

We tested whether thermal priming was sufficient to promote fitness metrics (photosynthetic efficiency [F_v_/F_m_], red channel intensity, growth rates), and whether these patterns were consistent between congeneric *Orbicella spp*. Overall, DTV-primed fragments exhibited significantly higher F_v_/F_m_ relative to stable-primed corals in *O. franksi* (*p* < 0.001), but not *O. faveolata* (Figure S3). Though F_v_/F_m_ of all DTV-primed fragments trended higher at all timepoints compared to stable-primed fragments for both coral species, only T_0_ and T_30_ timepoints were significantly higher in DTV-primed *O. franksi* and only T_30_ for *O. faveolata* (*p* < 0.05; Figure S3). Together, these patterns suggest that DTV priming had an initial impact on F_v_/F_m_ for both species, but these effects were transient and primed treatments did not differ at T_90_ (Figure 2A). Following heat challenge, all corals experienced reduced F_v_/F_m_ regardless of thermal priming treatment (*p* < 0.05; Figure 2A), except with stable-primed *O. franksi* where reductions in F_v_/F_m_ were observed under heat challenge, but differences were not significant (Figure 2A).

**Figure 2.**
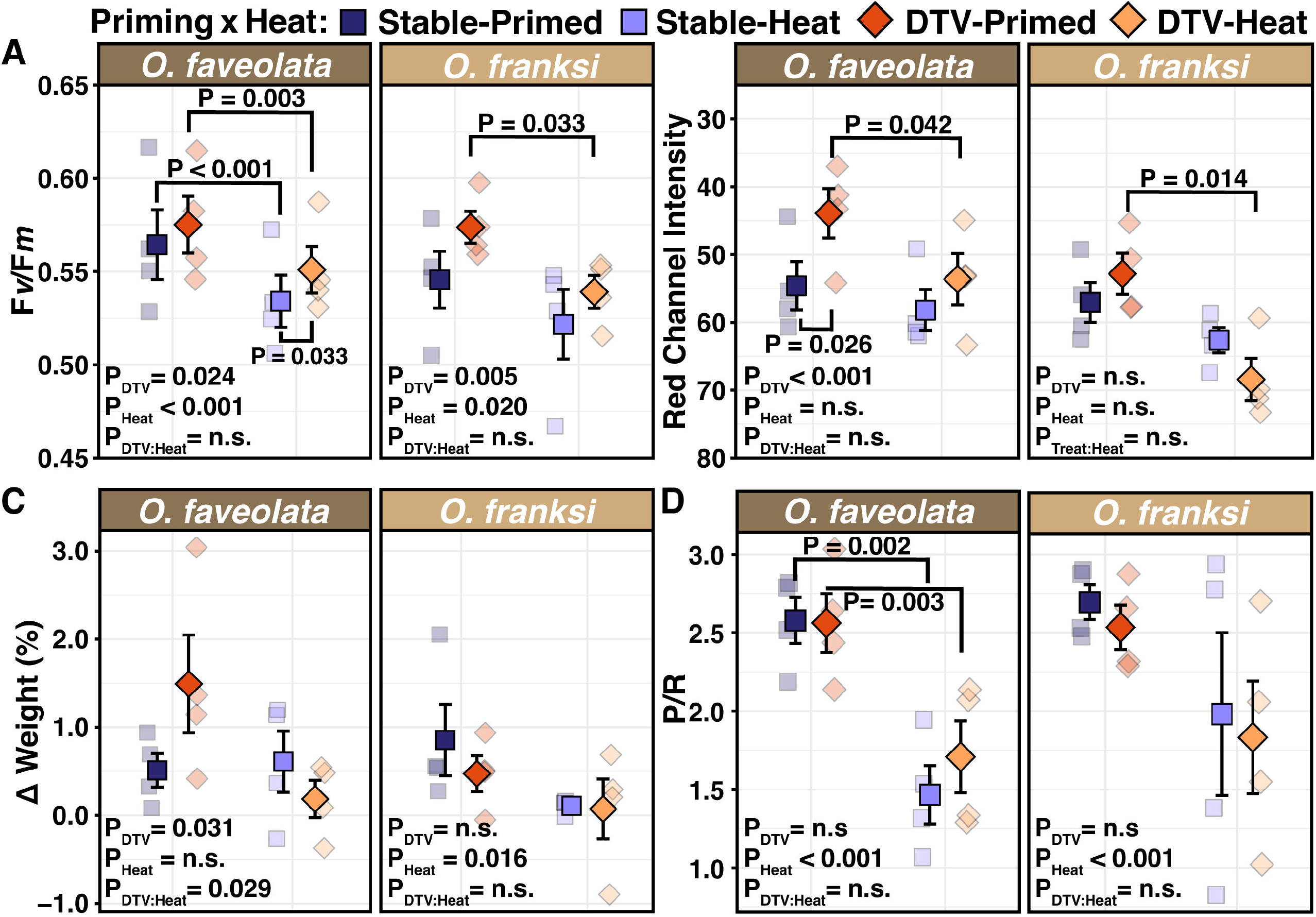
Coral physiology through thermal priming and heat challenge. Mean **(A)** photosynthetic efficiency (F*_v_*/F*_m_*), **(B)** mean red channel intensity (inverted to indicate bleaching: higher values = darker corals), **(C)** percent change in weight, and **(D)** Ratio of gross photosynthesis to dark respiration (P/R) rates quantified after thermal priming (left, T_90_) and heat challenge (right, T_119_) for *Orbicella faveolata* and *O. franksi.* Squares indicate stable thermal priming while diamonds indicate DTV priming. Treatments are colored as follows: Stable- Primed (blue), DTV-Primed (orange), Stable-Heat (light blue), DTV-Heat (light orange). Central points with error bars represent mean values ± standard error and transparent points represent values of individual fragments. *P*-values within each facet indicate results from *lmer*. Significant pairwise contrasts between time points (priming, heat challenge) are indicated by brackets above points, while pairwise contrasts within time points are indicated by brackets below points.

Red channel intensity (coral pigmentation) over the 90-day thermal priming showed differences through time for both host species, which were driven by lower pigmentation at T_30_ before light levels were lowered (*p* < 0.05; Figure S4). At the end of thermal priming, only DTV- primed *O. faveolata* were significantly darker than stable-primed fragments (*p* = 0.026, Figure 2B); however, only DTV-primed fragments exhibited significant paling under heat challenge, suggesting that DTV priming actually led to increased bleaching (*O. faveolata*: *p* = 0.042; *O. franksi*: *p* = 0.014; Figure 2B).

During thermal priming, *O. faveolata* and *O. franksi* both exhibited reduced growth over time (*p* < 0.05); however, only *O. faveolata* showed an effect of thermal priming (*p* = 0.031; Figure 2C) with higher growth in DTV-primed corals from T_60_-T_90_ (Figure S5). Importantly, this higher growth in DTV-primed fragments was not maintained under heat challenge (Figure 2C) and pairwise comparisons were not significant. Heat challenge only significantly reduced growth in *O. franksi*, regardless of priming treatment (*p* = 0.016; Figure 2C). Together, these patterns suggest that DTV priming had little effect on coral growth.

Oxygen flux was only measured at the end of the thermal priming experiment and following heat challenge. Thermal priming and heat challenge had no effect on gross photosynthesis for either species (Figure S6A). Heat challenge led to significantly higher dark respiration rates for *O. faveolata* (*p* = 0.02); however, no other effects of thermal priming or heat challenge were observed (Figure S6B). In contrast, heat challenge led to lower gross photosynthesis and dark respiration rate ratios (P/R) in both host species (*p* < 0.001 for both), but this reduction was much more pronounced in both thermal priming treatments in *O. faveolata* (Figure 2D). Overall, respiration and gross photosynthesis data (Figure 2D, S6) indicate that 90 days of DTV priming failed to modulate host or symbiont responses to subsequent heat challenge.

### Minimal influence of DTV and heat challenge on Symbiodiniaceae and bacterial communities

Following thermal priming, most fragments across both coral species hosted Symbiodiniaceae types belonging to *Durusdinium* type D1, putatively *D. trenchii* (Hume et al., 2020) (Figure 3). One *O. franksi* genotype (KF) hosted similar relative abundances of both *Breviolum minutum* (ITS2 type B1) and *D. trenchii* (Figure 3B, S2C). Background-levels, *i.e.* < 0.1 relative abundance per sample, of other Symbiodiniaceae types included *Breviolum* B2 and B1/B2 (Figure 3, S2C). Following heat challenge, three samples were dominated by *Symbiodinium* type A3 (Figure 3, S2C). Overall, ITS2-based results were similar to Symbiodiniaceae proportions observed from TagSeq reads; however, in the TagSeq analysis few *Symbiodinium* reads were observed and no samples were dominated by this genus (Figure S2). GLMM analysis indicated that thermal priming did not alter Symbiodiniaceae proportions, and no differences were detected between host species or in the interaction between DTV priming and heat challenge. There was some evidence of Symbiodinaceae compositional changes after heat challenge (*p* < 0.05), likely reflecting an increased relative abundance of *Symbiodinium* type A3 and *Breviolum* type B2 (Figure 3, S2C).

**Figure 3.**
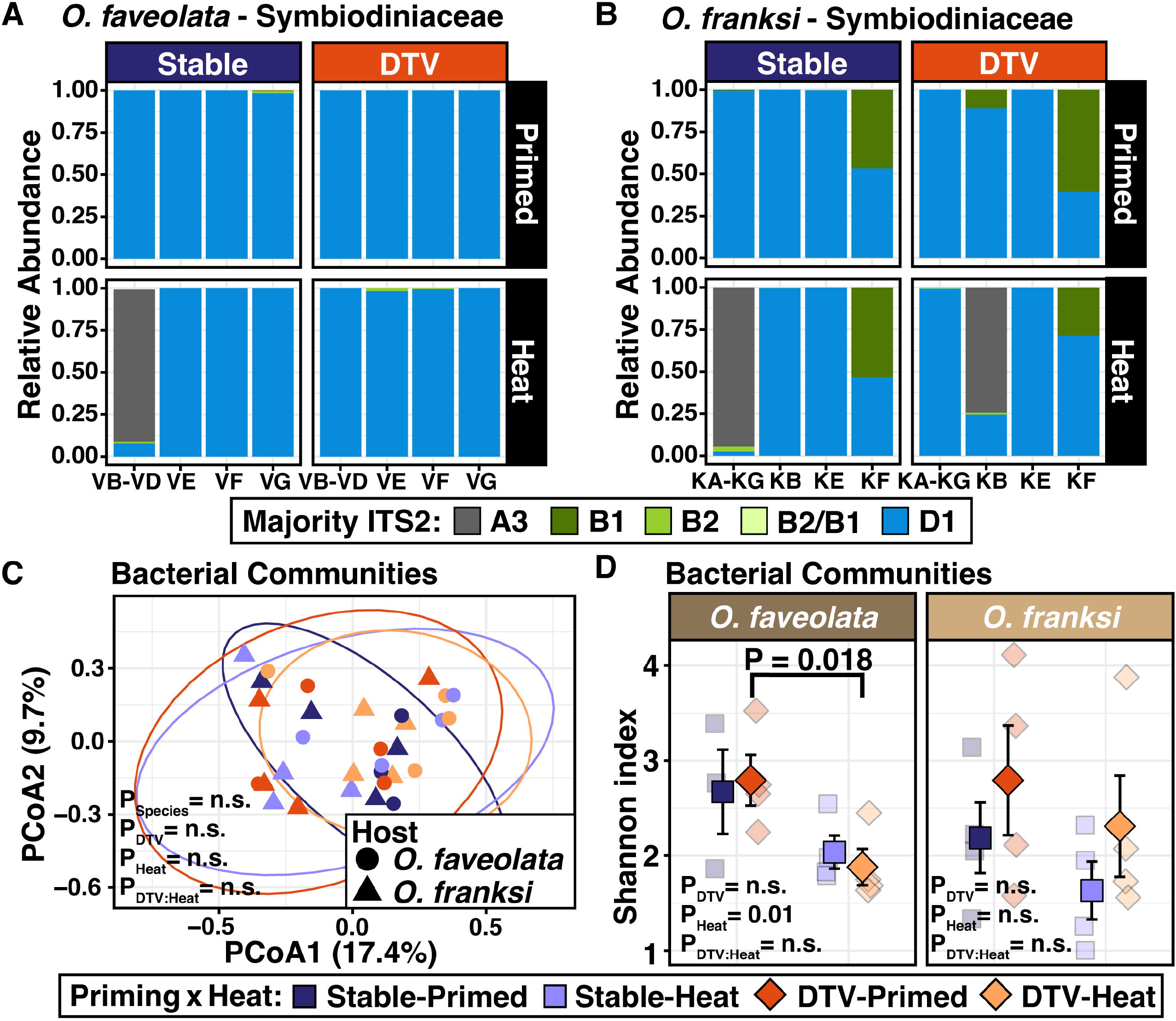
Symbiodiniaceae ITS2 type relative abundances and bacterial community diversity. Relative abundances of Symbiodiniaceae by ITS2 type for each **(A)** *Orbicella faveolata* and **(B)** *O. franksi* fragments (bars). Symbiodiniaceae types are shown as majority of each type profile (“Majority ITS2”), arranged by Stable-Primed (left) or DTV-Primed (right) during thermal priming (Primed, T_90_, top) or heat challenge (Heat, T_119_, bottom). **(C)** Principal coordinates analysis of Bray-Curtis dissimilarity of bacterial communities between samples’ relative abundances, where colors indicate experimental treatments and shapes denote host species. Ellipses are 95% confidence intervals. **(D)** Mean bacterial diversity estimated via Shannon index, compared between thermal priming and heat challenge. Central points with error bars represent mean values ± standard error.

Bacterial communities were largely composed of phyla Bacteroidota and Proteobacteria, with the most prominent families including Amoebophilaceae, Flavobacteriaceae, Teraskiellaceae, and Rhodobacteriaceae (Figure S7). GLMM analysis showed that bacterial community compositions did not differ by host species, thermal priming, or heat challenge (Figure 3C). Heat challenge reduced diversity of *O. faveolata* bacterial communities (*p* < 0.05), but thermal priming had no influence (Figure 3D). No differences in *O. franksi* bacterial community diversity were observed under DTV priming or heat challenge (Figure 3D). The same patterns were observed when examining Simpon’s index and when both metrics were calculated using rarefied 16S data, indicating that read depth differences did not affect these results.

### Little influence of DTV priming on host gene expression

Overall host gene expression (GE) profiles were significantly different between coral species (*p* < 0.001) and pre- and post- heat challenge (*p* < 0.05), but thermal priming had no influence (Figure 4A). No differences in host GE were detected for any thermal priming and heat challenge interactions (Figure 4B). For host GE, PC1 explains 15.2% of the variance among samples and showed separation by coral species, whereas PC2 (10.1% of variance) reflected GE variation due to heat challenge (Figure 4A-B). Numbers of host DEGs were similar across stable- and DTV- primed samples relative to heat challenge (stable: up 184, down 116; DTV: up 78, down 71; Figure S9).

**Figure 4.**
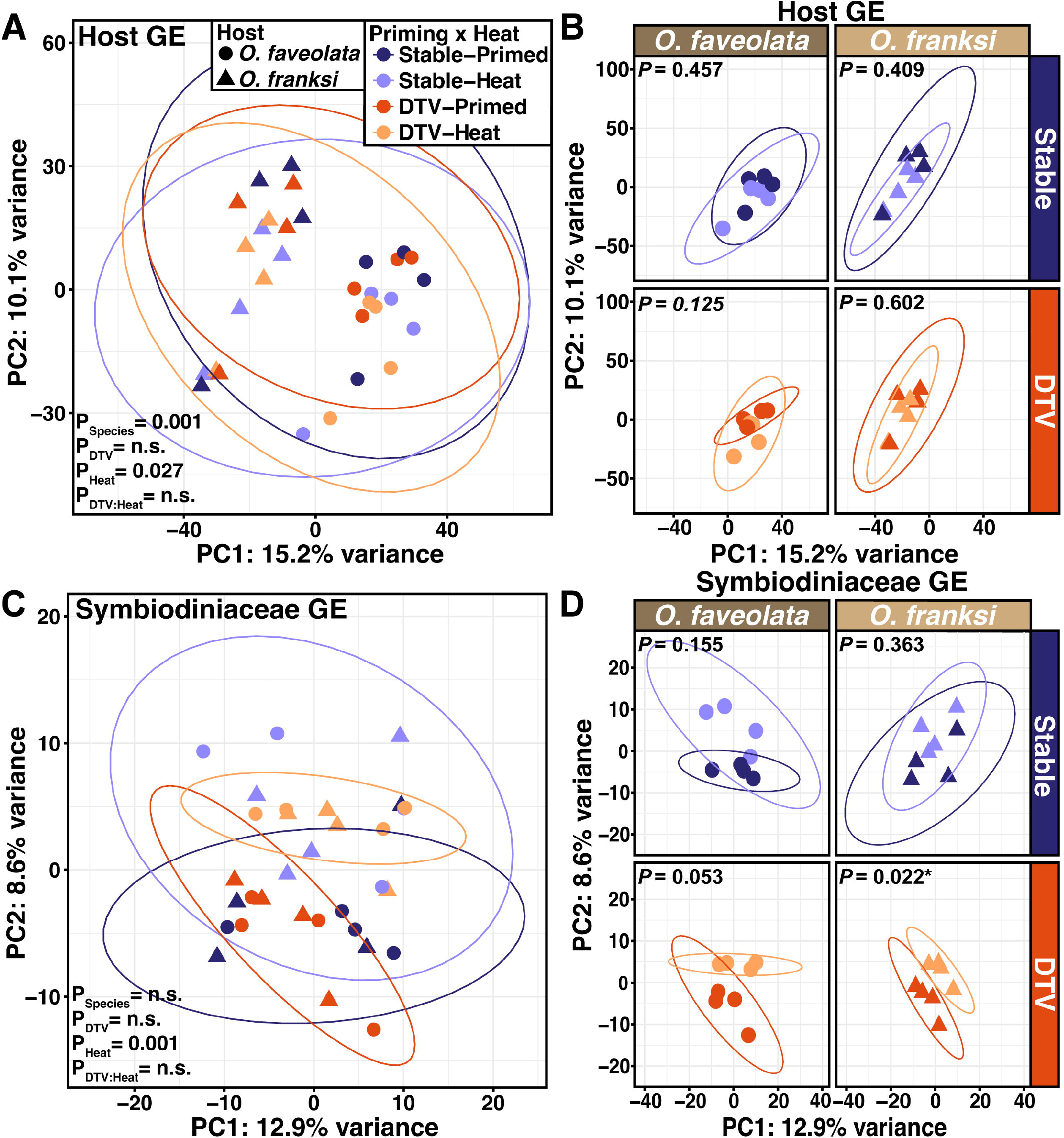
Principal component analysis (PCA) of coral host and symbiont gene expression profiles across DTV priming and heat challenge. **A)** Overall host gene expression profiles by host species (Species) and experimental treatments (Stable-Primed, DTV-Primed, Stable-Heat, DTV-Heat). **B)** Host gene expression profiles faceted by coral host (*O. faveolata*, *O. franksi*) and thermal priming treatment, **C)** Overall symbiont gene expression profiles by host species and experimental treatments. **D)** Symbiont gene expression profiles faceted by coral host (*O. faveolata*, *O. franksi*) and thermal priming treatments. Ellipses represent 95% confidence intervals. Significance was determined by PERMANOVAs (A,C) or pairwisePERMANOVA (B,C) for differences across treatments.

Weighted gene co-expression network (WGCNA) assigned 16,447 genes to 15 gene modules for the combined host analysis and 14 gene modules for host-specific analyses (*O. faveolata*, *O. franksi*). Module eigengene values were correlated with experimental treatments (DTV-Primed, Stable-Primed, DTV-Heat, Stable-Heat) and host traits (F_v_/F_m_, red channel intensity, buoyant weights, gross photosynthesis, dark respiration, and P/R; Figure 5, S10) for thermal primed (T_90_) and heat challenge (T_119_) time points. The combined host analysis indicated three modules correlating with DTV priming, the ‘cyan’ (705 genes), ‘black’ (1,190 genes), and ‘brown’ (1,400 genes) modules (Figure S10). The ‘cyan’ module was negatively correlated with DTV priming, while the ‘black’ module was negatively correlated with Stable-Heat, and the ‘brown’ module was negatively correlated with heat challenge for the combined host analysis (Figure S10). Since these modules were generally associated with thermal challenge and not unique to DTV priming, separate host-specific analyses were conducted. For *O. faveolata*, eigengene expression of the ‘brown’ (1504 genes) and ‘saddlebrown’ (218 genes) modules were positively correlated with DTV priming, while the ‘darkturquoise’ module (2,335 genes) was positively correlated with DTV-primed corals under heat challenge (Figure 5). The only modules significantly correlated with *O. franksi* treatments were the ‘brown’ (1,545 genes), ‘darkgrey’ (687 genes), and ‘ivory’ (682 genes) modules, with the ‘brown’ and ‘dark grey’ modules positively correlating with stable-primed corals and the ‘ivory’ module positively correlating with stable-primed corals under heat challenge (Figure 5).

**Figure 5.**
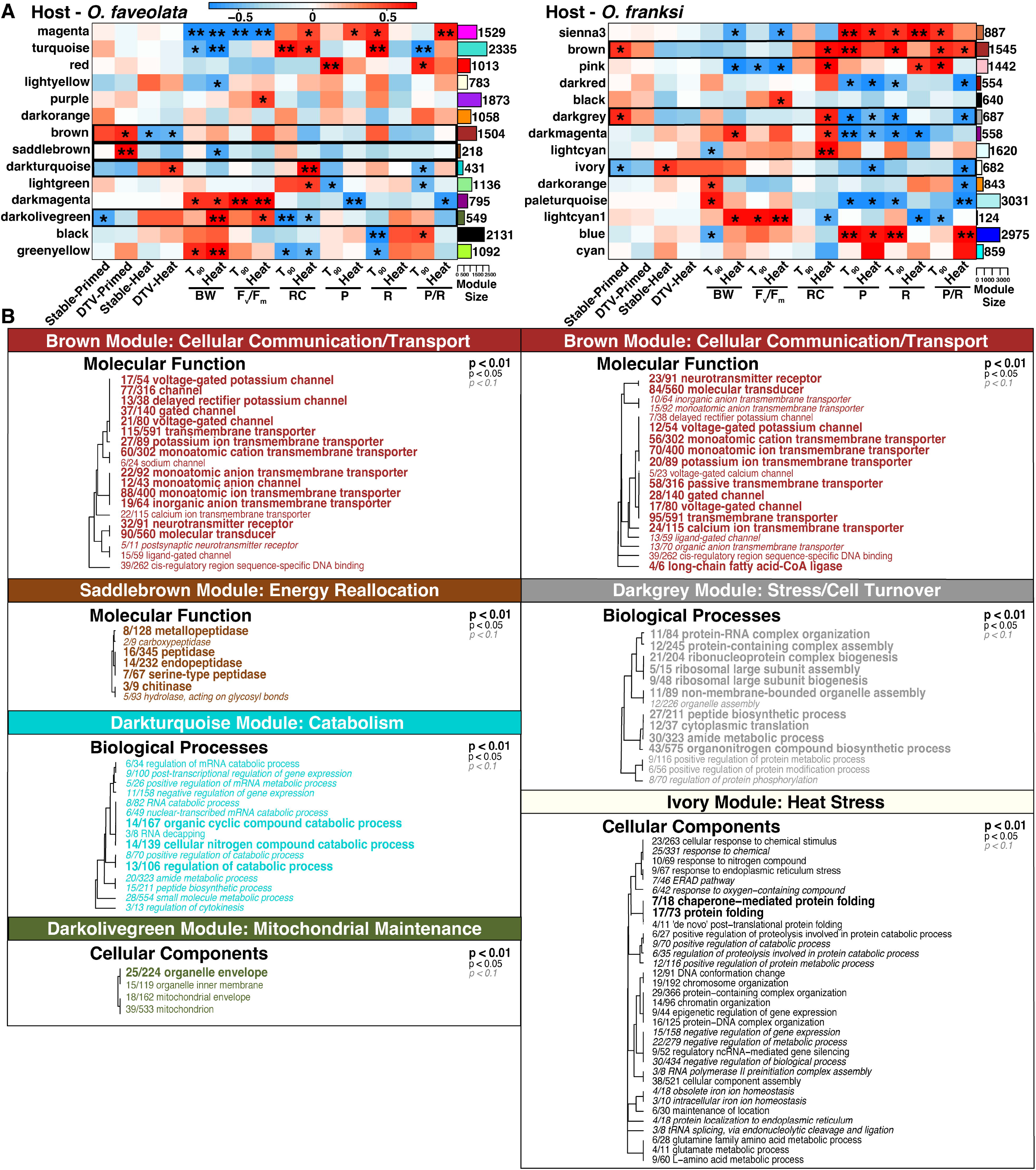
Weighted gene co-expression network analysis (WGCNA) of host module-trait relationships and associated module Gene Ontology (GO) enrichment. A) Module eigengene trait relationships identified by independent WGCNAs for *O. faveolata* (left) and *O. franksi* (right) host gene expression. Traits include all treatment conditions, change in buoyant weight (BW), F_v_/F_m_, Red Channel (RC), gross photosynthesis (P), dark respiration (R), and the ratio of gross photosynthesis to dark respiration (P/R). Correlation values ranging from 1 (red) to -1 (blue) and associated significant *p*-values (*p* < 0.05 indicated with a * and *p* < 0.001 indicated with **) are included within heatmap cells. B) Enriched GO terms of modules whose eigengene expression significantly correlated with treatments (Stable-Primed, DTV-Primed, Stable-Heat, DTV-Heat) for each host species are shown by molecular function (MF), cellular components (CC), and biological process (BP) categories. Font size and italics indicate levels of statistical significance as indicated by the inset key. Dendrograms show sharing of genes between GO terms and the fractions indicate the proportion of genes contained within the module relative to the total number of genes assigned to that GO term in the whole dataset.

GO enrichment analysis on the combined host WGCNA modules with high correlation to thermal priming treatments showed that the ‘cyan’ module had terms associated with histone binding (GO:0042393) and chromatin binding (GO:0003682), which were underrepresented in DTV-primed corals. The ‘black’ module showed underrepresentation of rRNA binding (GO:0019843), cytosolic ribosome (GO:0022626), and organonitrogen compound biosynthetic process (GO:1901566) in stable-primed corals. Lastly, the ‘brown’ module was underrepresented for cell-cell signaling (GO:0007267), synapse (GO:0045202), and transmembrane transporters (GO:0022857) in heat-challenged corals (Figure S10). Overall, the associations observed suggest classic heat stress responses, including the reduction of energetically costly cellular exchanges and communication during heat challenge in both host species.

In *O. faveolata,* WGCNA identified a ‘brown’ module that showed enrichment for transmembrane transportation and signaling genes, including terms such as potassium ion transport (GO:0015079; GO:0005267), voltage-gated channel activity (GO:0022843; GO:0005244; GO:0022832), and cell-cell signaling (GO:0007267). These pathways were thus overrepresented in DTV-primed corals but underrepresented under heat challenge (Figure 5). The *‘*saddlebrown’ module was enriched for damage control and recycling mechanisms and included proteolysis (GO:0006508) and peptidases (GO:0008233), indicating that DTV-primed corals were enriched for these pathways. In contrast, the ‘darkturquoise’ module was positively correlated with DTV-primed corals under heat challenge and and this module was enriched for metabolism and catabolism processes, such as cellular nitrogen compound catabolic process (GO:0046700; GO:0034655; GO:0044270) and amide metabolic process (GO:0043603; GO:0006518). Lastly, the ‘darkolivegreen’ *O. faveolata* module (549 genes) was negatively correlated with stable-primed corals and this module was enriched for terms associated with organelle envelopes (GO:0031967; GO:0031975).

In *O. franksi,* WGCNA analysis showed that all module eigengenes of interest positively (‘brown’ and ‘darkgrey’ modules) or negatively (‘ivory’) correlated with stable-primed corals (Figure 5). The *O. franksi ‘*brown’ module showed consistent patterns to the *O. faveolata* ‘brown’ module (Figure 5, S11), exhibiting similar enrichment of transmembrane transport (GO:0055085), cell-cell signaling (GO:0007267), and neurotransmitter receptor (GO:0042165; GO:0030594). The *O. franksi* ‘darkgrey’ module showed underrepresentation of cell component turnover, including rRNA binding (GO:0019843) and cytosolic ribosome (GO:0022626) in heat challenged corals. The ‘paleturquoise’ *O. franksi* module was not correlated with any treatments but expression of genes in this module were negatively correlated with oxygen flux measures (P, R, P/R; Figure S12) and included enrichment of proton-transporting ATPase complex (GO:0016469), which include V- and H-type ATPases that have been linked to symbiosome acidification (Barott et al., 2015).

The ‘brown’ modules for both host species showed similar patterns of enrichment and were both negatively correlated with heat challenge. Module comparisons showed high correlation between GO enrichment values (*τ* = 0.456, *p* < 0.001; Figure S11) and high overall semantic similarity (Figure S11). Half of the significantly enriched GO terms (10/20 terms) in these ‘brown’ modules were shared by both host species (Figure S11), suggesting that this module represents a conserved heat challenge response across these species.

### DTV priming led to stronger gene expression responses in Symbiodiniaceae under heat challenge

Overall Symbiodiniaceae GE profiles were influenced by heat challenge (*p* < 0.001, PERMANOVA), but not host species or DTV priming (Figure 4C) when data was not subset by host species. Interestingly, DTV-primed symbionts exhibited stronger GE differences in response to heat challenge than stable-primed symbionts; however, this difference was only significant for *O. franksi* (*p =* 0.22; Figure 4D) with *O. faveolata* showing marginal significance (*p* = 0.053; Figure 4D) when data were subset by host species. Tighter clustering of symbiont GE was also observed in both host species for DTV-primed symbionts following priming and heat challenge relative to stable-primed symbionts (Figure 4C). Similar to the host GE analysis, PC2 of the symbiont PCA (8.6% of variance) showed differences in GE due to heat challenge; however, no obvious sample clustering was observed along PC1 (12.9% of variance). Numbers of Symbiodiniaceae DEGs were similar between DTV- and stable-primed symbionts (DTV: up 9, down 25; stable: up 5, down 12; Figure S9). Mean Symbiodiniaceae GE plasticity did not differ between treatments; however, variance in plasticity differed (Levene: *p* < 0.05) or was marginally significant (Fligner’s: *p* = 0.056) among treatment groups with DTV-primed symbionts showing less variation in plasticity following heat challenge relative to stable-primed symbionts (pairwise Levene tests, *p* < 0.01; pairwise Fligner tests, *p* < 0.05; Figure S8).

## DISCUSSION

### Limited effects of DTV priming on coral physiology, symbioses, and gene expression

We tested whether diel thermal variation (DTV) modulates coral physiology, symbioses, and GE in *O. faveolata* and *O. franksi* from the FGB. After 90 days, DTV priming influenced select Symbiodiniaceae-related traits: DTV-primed corals exhibited higher photosynthetic efficiency (F_v_/F_m_) relative to stable-primed corals and *O. faveolata* were darker, suggesting greater pigment accumulation. Higher F_v_/F_m_ in DTV-primed corals may reflect increased photosystem recovery during nighttime reprieve (Klein et al., 2019; Putnam et al., 2010); however, DTV failed to measurably alter any other physiological traits including photosynthetic rates or growth in either host species. While increases in F_v_/F_m_ and growth under DTV have been reported in other coral taxa (Aichelman et al., 2025; Bay & Palumbi, 2015; DeMerlis et al., 2025; Putnam et al., 2010), neutral and negative responses have also been observed (DeMerlis et al., 2025; Putnam et al., 2010; Putnam & Edmunds, 2011; Schoepf et al., 2022). Variation among studies may reflect differences in experimental duration (Grottoli et al., 2021), scale of DTV, acclimation time, or life history and thermal niche differences (Darling et al., 2012; Silbiger et al., 2019). Regardless, our results indicate that modest increases in symbiont-related traits in DTV-primed corals failed to translate into enhanced metabolic performance or host fitness.

We next examined how DTV priming influenced symbiotic associations. Most Symbiodiniaceae communities were dominated by *D. trenchii* regardless of treatment, contrasting prior reports of *B. minutum* dominance in *Orbicella* spp. from FGB (Green et al., 2014). *Orbicella* host diverse Symbiodiniaceae across the Caribbean (Kemp et al., 2015; Thornhill et al., 2006) and can shift toward *D. trenchii* dominance under aquarium conditions (Gantt et al., 2023, 2024), which was also previously observed for these same FGB genotypes in a separate study (Strader et al., 2024). Like Symbiodiniaceae communities, bacterial community composition and diversity was stable across treatments after thermal priming (Figure 3C). This stability may reflect *ex situ* conditions lacking novel microbial inputs (Carrier & Reitzel, 2017) or an inherently less responsive microbiome in *Orbicella* spp. Although FGB coral microbiomes remain uncharacterized (but see sponge (Shore et al., 2021) and water column (Doyle et al., 2022) descriptions), similar microbiome resistance has been observed in *O. franksi* in Panamá (Prada et al., 2022) and *O. faveolata* in Puerto Rico (Tracy et al., 2015), consistent with other microbiome “regulator” corals (Ziegler et al., 2019). Overall, we found no support for shifts in Symbiodiniaceae or bacterial communities under DTV conditions.

GE patterns can predict responses to thermal stress (Bellantuono et al., 2012; Dixon et al., 2015) and front-loading of certain genes can increase coral stress resistance (Barshis et al., 2013; Palumbi et al., 2014). DTV priming could front-load gene pathways that increase thermal resistance; however, we observed little GE response to DTV priming for either coral species. In *O. faveolata*, DTV priming was positively associated with modules enriched for cellular communication/transport (‘brown’ module) and energy reallocation (‘saddlebrown’ module), suggesting these functions are involved in long-term DTV acclimation. In contrast, no modules were correlated with DTV priming in *O. franksi*. ‘Brown’ modules in both host species were highly homologous, but positively correlated with opposite thermal priming treatments, with cellular communication/transport enriched in DTV-primed *O. faveolata* and stable-primed *O. franksi.* In *O. franksi*, this ‘brown’ module was positively associated with symbiont-linked traits (red channel, P, R, P/R), suggesting host regulation of symbiont performance or stability within the host. Overall, these host GE results suggest muted, although somewhat species-specific, responses to DTV priming. Species-specific responses to novel environments have previously been observed within the *Orbicella* complex, with *O. franksi* exhibiting greater physiological plasticity (Prada et al., 2022). While our data do not support large plasticity differences between species, it is possible that *Orbicella* spp. exhibit rapid acclimation (Bay & Palumbi, 2015) and we therefore missed the critical window for observing GE plasticity in response to DTV priming (Rivera et al., 2021).

While no shifts in Symbiodiniaceae communities were observed in response to DTV, *D. trenchii* GE supports functional responses to DTV priming. Across host species, DTV-primed Symbiodiniaceae exhibited more similar GE profiles after priming and heat challenge, and these patterns were more variable in stable-primed symbionts. As discussed above, DTV-primed corals also exhibited enhanced photochemical efficiency. Together, these patterns observed in DTV- primed Symbiodiniaceae suggest that priming promotes a highly coordinated acclimation response in *D. trenchii. D. trenchii* GE was not significantly influenced by host species, even though previous work on FGB *O. franksi* and *O. faveolata* hosting *Breviolum minutum* demonstrated clear host-specific patterns of symbiont expression (Strader et al., 2024). This paucity of host effects may reflect differences between homologous and heterologous symbiont pairings (Lust et al., 2025; Wuerz et al., 2023), or more broadly, between long co-evolved partnerships versus recently established associations, as *D. trenchii* is hypothesized to have been recently introduced to Atlantic corals (Pettay et al., 2015).

### *DTV does not promote heat tolerance in* Orbicella spp

DTV priming did not improve *Orbicella* thermal resistance, and both species exhibited comparable physiological declines during the heat challenge, regardless of prior DTV exposure. These responses contrast with studies linking DTV acclimation with reduced coral bleaching (Palumbi et al., 2014; Safaie et al., 2018), but corroborate others highlighting tradeoffs between environmental variability and thermal resistance (Klepac & Barshis, 2020; Schoepf et al., 2022). Although DTV-primed corals exhibited higher photosynthetic efficiency, this advantage did not translate into protection from bleaching-associated declines during the heat challenge. Resilience, *i.e.* recovery following disturbance, is an alternative to stress resistance, and both strategies are reported in natural coral populations (Lam et al., 2020). While we did not assess post-stress recovery dynamics, a recent study with *Siderastrea siderea* similarly showed that earlier impacts of DTV were not sustained through a heat challenge and failed to improve health metrics during a recovery period (Aichelman et al., 2025). While additional work examining longer-term acclimation strategies across other coral taxa is needed, our findings suggest that DTV alone may not reliably enhance coral resistance to thermal stress.

Despite physiological indicators of coral bleaching, Symbiodiniaceae and bacterial communities remained mostly stable after the heat challenge. This result is notable given that heat stress and other environmental changes tend to restructure both algal symbiont and bacterial associations across diverse coral taxa (A. C. Baker, 2003; McDevitt-Irwin et al., 2017) (but see (Goulet, 2006)), including *O. faveolata*-associated Symbiodiniaceae (Cunning et al., 2015). The absence of major Symbiodiniaceae shifts under thermal challenge may be attributed to the dominance of *D. trenchii*. Claar et al. (2020) observed that colonies previously hosting *Durusdinium* did not shuffle symbiont communities under heat stress, while colonies hosting *Cladocopium* shuffled to *Durusdinium*. In our study, bacterial communities of both coral species were unresponsive to the heat challenge, except *O. faveolata* showed decreased diversity. While some studies report increased bacterial alpha diversity in response to thermal stress, presumed to be a disruption of “stable” microbiome functioning (McDevitt-Irwin et al., 2017), findings from *O. faveolata* in Mexico showed stable bacterial diversity between control and heat stressed treatments (Avila-Magaña et al., 2021). This study also found that bacterial communities were less metabolically active under heat stress than the coral host and Symbiodinaceae, suggesting bacterial communities played a less important role in the heat stress response. Our results provide support for remarkably stable bacterial and algal communities within *Orbicella* under mesocosm-based heat challenge; however, we cannot rule out functional responses of these communities that were not evaluated here.

Despite strong heat-driven transcriptional responses, unique responses of DTV-primed corals to heat challenge were not clear. All combined host WGCNA modules correlating with experimental treatments were linked to heat challenge rather than DTV priming, encompassing cell signaling, ribosomal protein biosynthesis, and chromatin/post-transcriptional control with the latter associated with declines in photosynthetic efficiency and indicative of a tradeoff in post-transcriptional activity (Seneca & Palumbi, 2015). Similarly, separate host WGCNAs revealed a conserved heat response, marked by reduced cellular signaling and transport. In *O. faveolata*, heat stress was further characterized by enrichment of catabolic pathways and mitochondrial maintenance genes (Davies et al., 2016), which were positively correlated with reduced red channel intensity, F_v_/F_m_, and growth rates. In contrast, *O. franksi* was associated with modules enriched for canonical stress-response including cell turnover and protein misfolding (Dixon et al., 2020; Ruiz-Jones & Palumbi, 2017). It is worth noting that DTV- Primed *O. franksi* exhibited a more muted transcriptional response, both following priming and heat challenge, compared to Stable-Primed *O. franksi*, which matches responses observed in DTV primed *Acropora* species (Bellantuono et al., 2012). This muted response suggests that DTV priming induces molecular preconditioning; however, as host GE was largely driven by thermal stress irrespective of priming treatment, there is little evidence that DTV priming led to GE changes that improved coral thermal resistance.

While DTV-Primed Symbiodiniaceae maintained reduced among-sample variation relative to Stable-Primed symbionts, significant clustering of overall GE was driven by heat challenge in both host species. Similarly DEGs were found between Symbiodiniaceae prior to and following heat challenge, with no DEGs were associated with DTV priming alone. This lack of Symbiodiniaceae GE response is not novel, as many studies have observed this pattern *in hospite* (Barshis et al., 2014; Davies et al., 2018; Leggat et al., 2011). While this paucity of response to stress by Symbiodiniaceae has been attributed to differences in transcriptomic machinery (Baumgarten et al., 2013), Aichelman et al. (2024) demonstrated GE patterns consistent with host buffering when comparing Symbiodiniaceae responses *in hospite* versus in culture, suggesting that hosts tightly regulate the symbiosome (Barott et al., 2015). Because coral hosts likewise showed no response to DTV priming, it remains unclear whether the Symbiodiniaceae’s lack of response arises from the same underlying mechanisms.

### Conclusions

Here, DTV exposure resulted in minimal physiological or GE plasticity in congeneric coral holobionts and their symbiont *D. trenchii*. Although DTV increased photosynthetic efficiency in both Orbicellids and pigmentation in *O. faveolata*, this DTV priming failed to facilitate thermal stress tolerance. Symbiodiniaceae and bacterial community compositions also remained stable after DTV priming in both coral species. These findings suggest mean temperature may play a stronger role in shaping *Orbicella* holobiont metrics than DTV, and that DTV alone does not mitigate bleaching susceptibility. Furthermore, given that temperature is just one of many diel variables on reefs, other factors such as water chemistry, light, nutrient availability, and turbidity demand greater attention. Incorporating these fluctuations will better represent natural conditions and will lead to a stronger understanding of the acclimatory responses of holobionts to future climate change.

## DATA AVAILABILITY STATEMENT

All raw sequencing data are available on NCBI’s Sequence Read Archive (ITS2: PRJNA1442140, 16S: PRJNA1442122, RNA: PRJNA1438202). Processed data files and scripts used in analyses are available at https://github.com/shegantt/DTV_Ofav_Ofra and are additionally hosted on Zenodo (DOI: Will update with a DOI after review and acceptance).

## ACKNOWLEDGEMENTS

We are grateful to the Flower Garden Banks National Marine Sanctuary, which allowed coral collections under permit FGBNMS-2018-006. Thank you to Brooke Benson, Brianne Dent, Isabela Trumble, and Kian Thompson for coral husbandry assistance, and James Fifer for assistance during experimentation. We thank the BU Marine Program and Justin Scace for access to the seawater facility. Thank you also to significant input on analyses and writing from Davies lab members present and past, Drs. Peter Buston, Jennifer Bhatnagar, and Jeffrey Marlow. We respectfully acknowledge that Boston University’s Charles River Campus stands on the ancestral and unceded homelands of the Massachusett (Massachusett, Pawtucket/Naumkeag) people and honor their continuing relationship with this land.

## AUTHOR CONTRIBUTIONS

SWD and NGK conceived of the study design, and SWD provided funding from Boston University start-up funds. NGK, HEA, and HER performed physiological and sampling protocols during the experiment. AKH and EM conducted laboratory-based molecular work. CED processed oxygen flux data. SEG and NGK conducted data analyses, created figures, and drafted the manuscript, with significant contributions by SWD. All authors provided manuscript revisions and final submission approval.

