## Supplemental Materials for "Diel thermal variability does little to mitigate bleaching in two endangered coral holobionts"

**Table S1. Water quality metrics (temperature, salinity, concentrations of nitrate, phosphate, calcium, alkalinity, and magnesium) compared between diel thermal variability (DTV)-Primed and Stable-Primed thermal priming treatments.** Mean values ± standard deviation are shown. Variance between treatments was compared via Levene's test and difference in means compared via Wilcoxon rank sum test for all variables, except for salinity, which met model assumptions and a one-way ANOVA was used ($\alpha = 0.05$).

| Metric (unit, # of measurements) | Stable (mean ± SD) | DTV (mean ± SD) | Variance comparison | Mean comparison |
|---|---|---|---|---|
| Temperature (˚C, $n$ = 18,170) | 25.67 (± 0.16) | 25.73 (± 1.36) | $F_{1,18159} = 22757$ $p < 2e\text{-}16$*** | W = 42774464 $p <$ 1.064$e$-05*** |
| Salinity (ppt, $n$ = 183) | 34.41 (± 0.87) | 34.31 (± 0.95) | n.s. | n.s. |
| Nitrate (ppm, $n$ = 21) | 0 (± 0) | 0 (± 0) | NA | NA |
| Phosphate (ppm, $n$ = 17) | 0.04 (± 0.05) | 0.03 (± 0.04) | n.s. | n.s. |
| Calcium (ppm, $n$ =21) | 433.10 (± 43.58) | 437.50 (± 27.41) | n.s. | n.s. |
| Alkalinity (dKH, $n$ = 21) | 8.47 (± 1.95) | 8.11 (± 1.84) | n.s. | n.s. |
| Magnesium (mg/L, $n$ = 21) | 1370.50 (± 61.66) | 1361.36 (± 44.27) | n.s. | n.s. |

**Table S2. Holobiont sequencing information through processing, trimming, mapping, and gene counts.** SampleIDs indicate coral species (K = *O. franksi*, V = *O. faveolata*), individual genotype (A, B, C, E, F, G), and fragment number. Included are sequence counts from unfiltered fastq files (Raw Reads) and files after trimming and filtering (Trimmed Reads) sequences. Host Counts and Symbiont Counts indicate reads mapping to the *O. faveolata* host genome and *D. trenchii* symbiont genome. Mapping Efficiency (%) is the percentage of total host and symbiont counts (Host Counts + Symbiont Counts) relative to the total number of filtered reads (Trimmed Reads). Host and Symbiont Genes indicate the number of putative genes with counts from host and symbiont mapping. *Genotypes KA and VB (asterisks) were removed because they were clonemates with KG and VD, respectively.

| SampleID | Raw Reads | Trimmed Reads | Host Counts | Symbiont Counts | Mapping Efficiency (%) | Host Genes | Symbiont Genes |
|---|---|---|---|---|---|---|---|
| KA1-1* | 5454393 | 1713114 | 915183 | 437430 | 79.0 | 15029 | 2037 |
| KA3-1* | 5705638 | 1566591 | 786852 | 546207 | 85.1 | 14454 | 1945 |
| KA4-2* | 6580487 | 1831662 | 963206 | 573632 | 83.9 | 15107 | 2015 |
| KA5-2* | 6929713 | 1806197 | 1286196 | 193257 | 81.9 | 15413 | 1404 |
| KB13-2 | 6665162 | 2258501 | 1066048 | 801905 | 82.7 | 15300 | 2205 |
| KB28-1 | 6513247 | 2364810 | 983062 | 1044171 | 85.7 | 15235 | 2266 |
| KB39-1 | 6276484 | 808749 | 452141 | 233697 | 84.8 | 12605 | 1541 |
| KB39-2 | 5905208 | 2027863 | 1207688 | 432257 | 80.9 | 15387 | 1972 |
| KE13-1 | 5744076 | 2066313 | 932055 | 825846 | 85.1 | 15227 | 1785 |
| KE14-1 | 5694559 | 1481334 | 1035177 | 146307 | 79.8 | 14821 | 1284 |
| KE31-2 | 8650934 | 1559284 | 1158441 | 105942 | 81.1 | 15139 | 1026 |
| KE9-2 | 6020069 | 1888099 | 1038914 | 570093 | 85.2 | 15320 | 2049 |
| KF19-1 | 6272629 | 1674313 | 1123245 | 126680 | 74.7 | 14935 | 1022 |
| KF19-2 | 8807209 | 2200829 | 1176579 | 535874 | 77.8 | 15102 | 1928 |
| KF23-2 | 9225466 | 1519045 | 1051869 | 84708 | 74.8 | 14630 | 661 |
| KF9-1 | 7930530 | 1109431 | 628007 | 157061 | 70.8 | 12889 | 1113 |
| KG12-1 | 7193077 | 2067895 | 1051820 | 689974 | 84.2 | 15025 | 2079 |
| KG12-2 | 7410684 | 1907147 | 1099529 | 506562 | 84.2 | 15013 | 1921 |
| KG31-1 | 5988279 | 1735758 | 1222731 | 191830 | 81.5 | 15395 | 1466 |
| KG31-2 | 5222728 | 1573192 | 900488 | 407233 | 83.1 | 14966 | 1884 |
| VB11-1* | 6573910 | 2031826 | 1286493 | 454456 | 85.7 | 15570 | 2159 |
| VB11-2* | 6368067 | 1122908 | 605945 | 358819 | 85.9 | 13981 | 1785 |
| VB1-2* | 6265451 | 1903223 | 984297 | 644044 | 85.6 | 15252 | 2063 |
| VB5-1* | 7461661 | 2347548 | 967810 | 929338 | 80.8 | 15277 | 2198 |

| | | | | | | | |
|---|---|---|---|---|---|---|---|
| VD10-1 | 6464677 | 1876021 | 1130491 | 433837 | 83.4 | 15342 | 1902 |
| VD15-2 | 7486510 | 2071516 | 1182197 | 565770 | 84.4 | 15464 | 2001 |
| VD17-1 | 8107613 | 1870435 | 984099 | 575633 | 83.4 | 15256 | 2004 |
| VD17-2 | 7697623 | 2185702 | 1504792 | 312162 | 83.1 | 15763 | 1772 |
| VE14-2 | 6851480 | 2143480 | 828850 | 1065778 | 88.4 | 14360 | 2131 |
| VE18-1 | 6671230 | 2183777 | 1374223 | 462595 | 84.1 | 15609 | 1961 |
| VE18-2 | 6937246 | 2071858 | 1406824 | 338973 | 84.3 | 15562 | 1795 |
| VE30-1 | 8010374 | 2500965 | 1019343 | 1169007 | 87.5 | 15128 | 2235 |
| VF23-2 | 7340100 | 2424200 | 1210168 | 912359 | 87.6 | 15480 | 2220 |
| VF31-1 | 8629965 | 2281800 | 979962 | 1039138 | 88.5 | 14708 | 2177 |
| VF40-1 | 8826452 | 1274918 | 811199 | 266919 | 84.6 | 13601 | 1548 |
| VF40-2 | 8850477 | 1116559 | 758409 | 200483 | 85.9 | 13531 | 1425 |
| VG1-1 | 7421010 | 1930161 | 1371576 | 261628 | 84.6 | 15541 | 1642 |
| VG1-2 | 7235264 | 1139463 | 743384 | 235589 | 85.9 | 14028 | 1557 |
| VG17-2 | 7173769 | 2575007 | 1089053 | 1134249 | 86.3 | 15486 | 2271 |
| VG6-1 | 7842604 | 144238 | 499116 | 456301 | 83.5 | 13535 | 1902 |

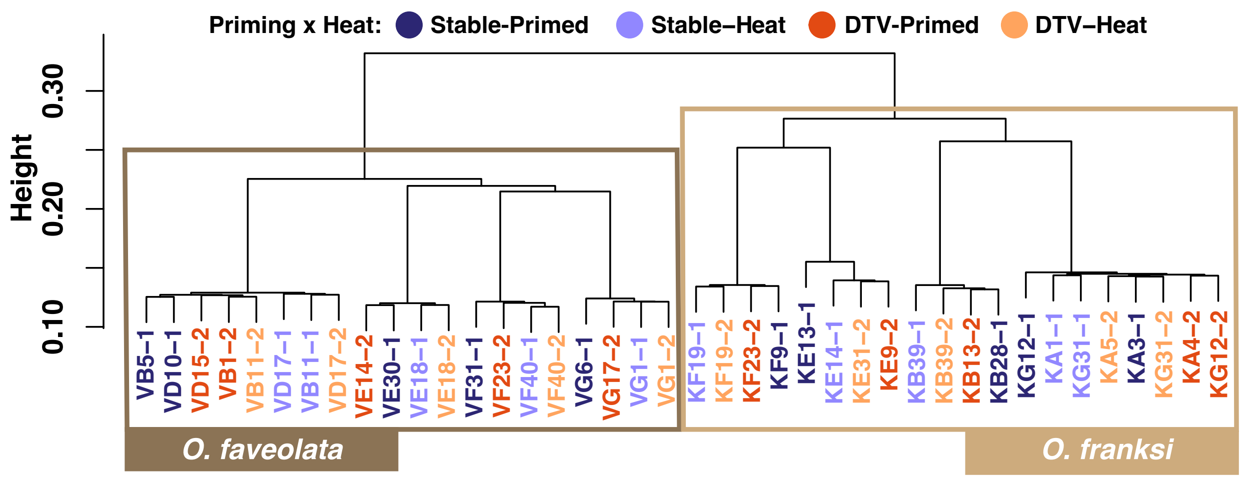


**Figure S1. Identity by state (IBS) cluster dendrogram of all samples using single nucleotide polymorphisms (SNPs) in TagSeq data.** Nodes below a height of 0.2 represent the cutoff for putative clonal groups where all fragments belong to the same genotype. Treatment conditions are indicated in text color and species are indicated with boxes colored as dark brown for *O. favoelata* and light brown for *O. franksi.* It is observed that colonies VB/VD and KG/KA are clonemates.

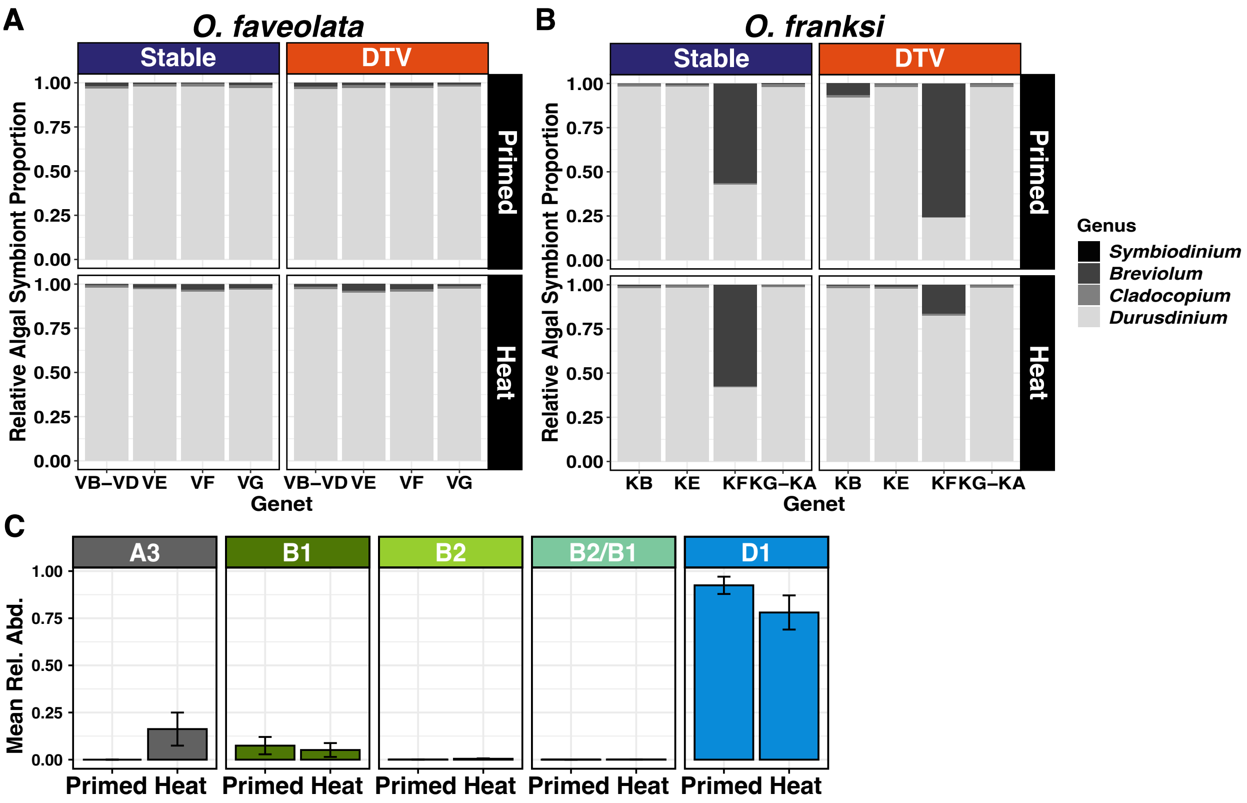


**Figure S2. Relative abundances of Symbiodiniaceae genera determined by single nucleotide polymorphisms (SNPs) in TagSeq data and mean relative abundances of each Symbiodiniaceae type from ITS2 sequencing.** Symbiodiniaceae genera relative abundances for each coral fragment by SNPs from TagSeq data for **(A)** *Orbicella faveolata* and **(B)** *O. franksi* from Stable (left) or DTV (right) thermal priming treatments and time points sampled post priming (Primed, top) or heat challenge (Heat, bottom). Each bar represents a single fragment with genotypes/clonemates indicated on the x-axis. **(C)** Mean relative abundances for majority ITS2 types from ITS2 sequencing. All samples, compared between the end of the DTV-priming ($T_{90}$, Primed) and heat challenge ($T_{119}$, Heat), where error bars denote standard error.

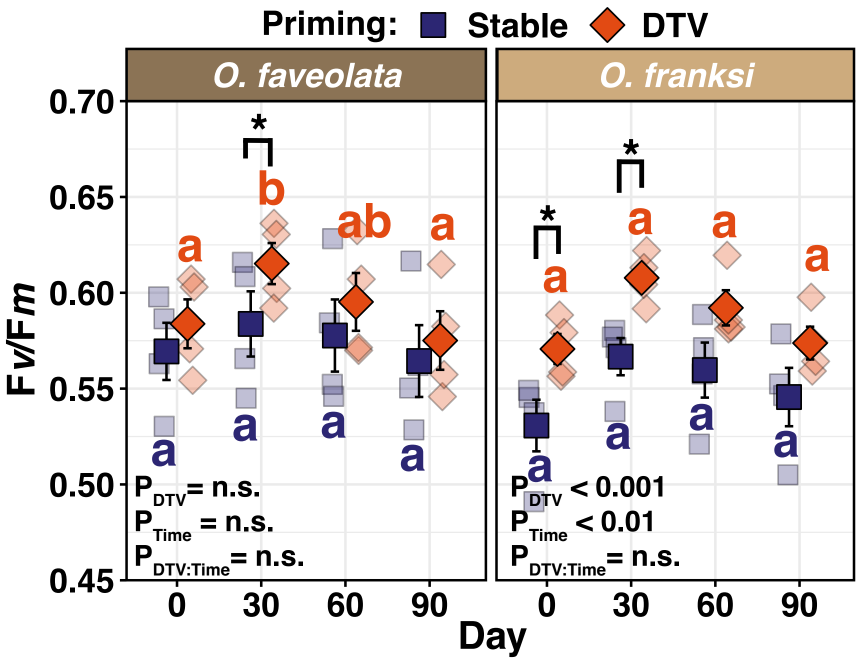


**Figure S3. Changes in $F_v/F_m$ through thermal priming treatment.** Mean $F_v/F_m$ +/- standard error (SE) of Stable-Primed and DTV-Primed *Orbicella faveolata* (left) and *O. franksi* (right) with each point representing the mean value of a fragment at that timepoint. Significance was determined by *lme* models within each coral species and pairwise significance values are shown by letters. Significance between Stable- and DTV-Primed corals within timepoints are indicated with black brackets and asterisks.

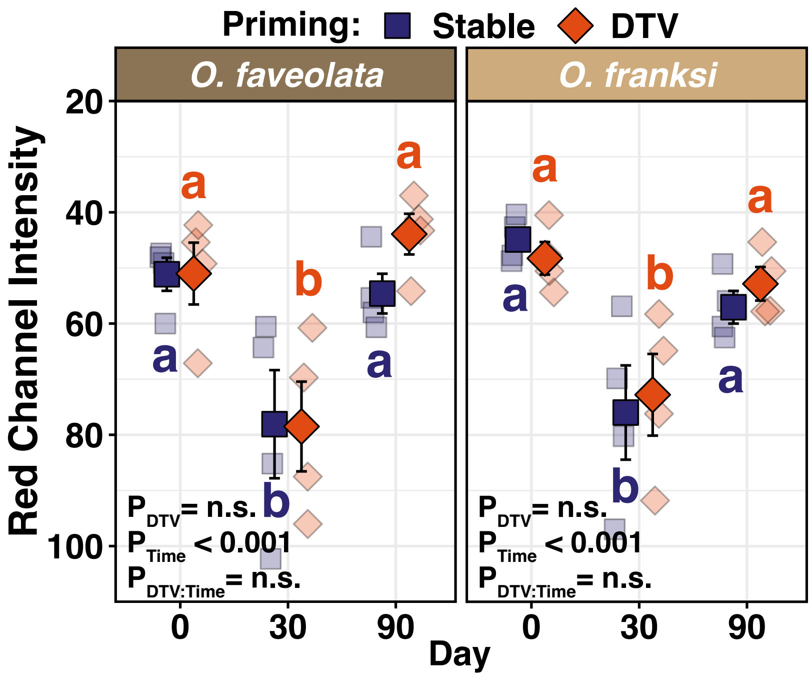


**Figure S4. Red channel intensity (proxy for coral bleaching) through thermal priming.** Mean red channel intensity +/- standard error (SE) of DTV-Primed and Stable-Primed *Orbicella faveolata* and *O. franksi* with each point representing the mean value of an individual at that timepoint. Red channel intensity values were inverted to more intuitively display a loss in color, *i.e.* paling, where higher values = darker pigmentation. Significance was determined by *lme* models within each coral species and pairwise significance values are shown by letters. Faded points behind means indicate the genotype means for each time point. No significant differences across treatment comparisons within timepoints were observed. No images were available for analysis from $T_{60}$ due to technical issues.

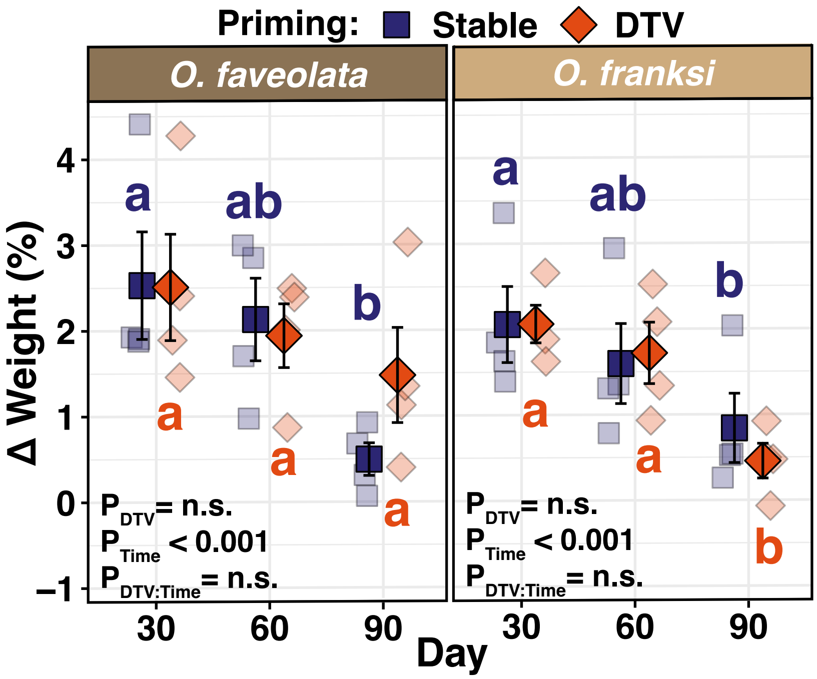


**Figure S5. Change in coral growth through thermal priming.** Mean growth +/- standard error (SE) of Stable-Primed and DTV-Primed *Orbicella faveolata* and *O. franksi* with each point representing the mean value of each genotype at that timepoint. Here, growth was measured as percent change in buoyant weight between time points (Day 30 = $T_{30}$-$T_0$, Day 60 = $T_{60}$–$T_{30}$, Day 90 = $T_{90}$-$T_{60}$). Significance was determined by *lme* models within each coral species and pairwise significance values are shown by letters. No significant differences across treatment comparisons within timepoints were observed.

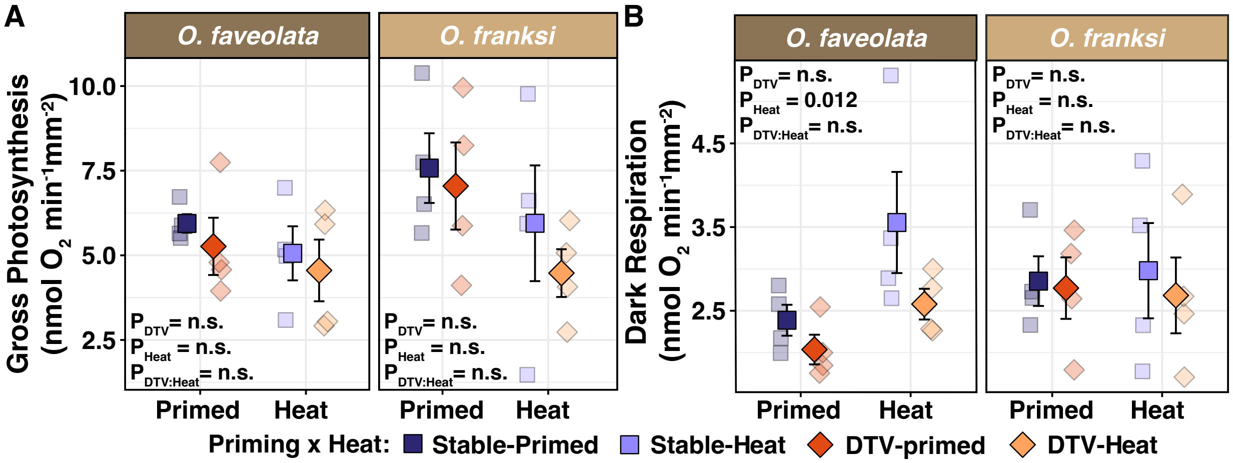


**Figure S6. Respirometry measures of A) Gross Photosynthesis and B) Dark Respiration following thermal priming and heat challenge.** Mean oxygen concentrations +/- standard error (SE) for gross photosynthesis and dark respiration of Stable-Primed and DTV-Primed *Orbicella faveolata* and *O. franksi* with each point representing the mean value of each genotype at that timepoint. Significance was determined by *lmer* models within each coral species and pairwise significance values are shown by letters. No significant differences were observed within or across treatment comparisons over the timepoints.

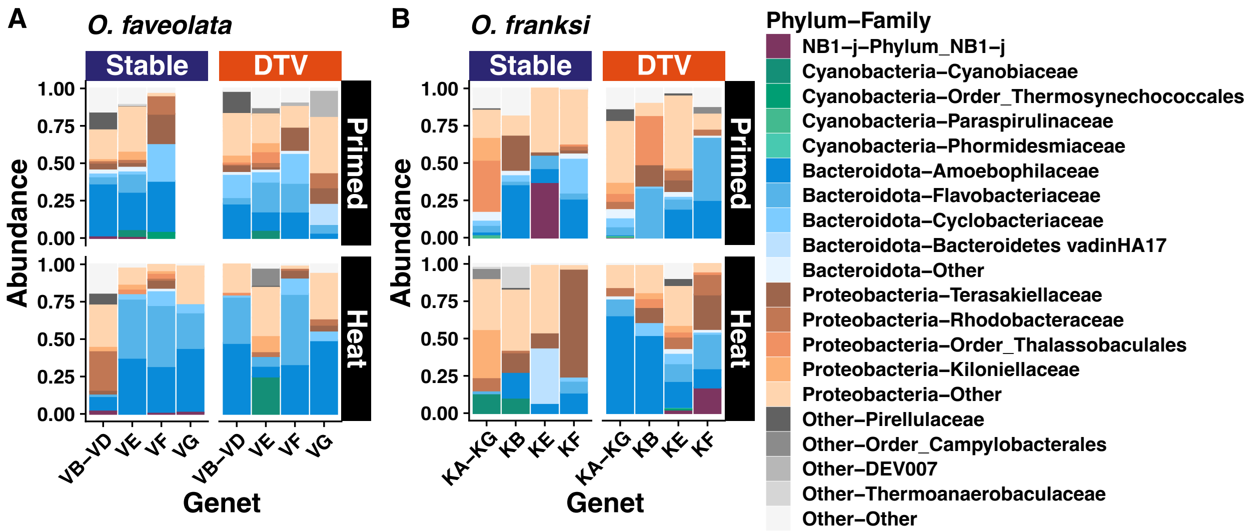


**Figure S7. 16S Bacterial communities from (A) *Orbicella faveolata* and (B) *O. franksi* following thermal priming and heat challenge.** Relative abundance of the total bacterial community compositions, where genotypes and thermal priming treatments (Stable-Primed or DTV-Primed) are arranged as columns and heat challenge treatment ("Primed" = $T_{90}$, "Heat" = $T_{119}$) as rows. Colors represent the top four most abundant bacterial phyla and top four families within each phyla, and all other taxa are labeled "Other."

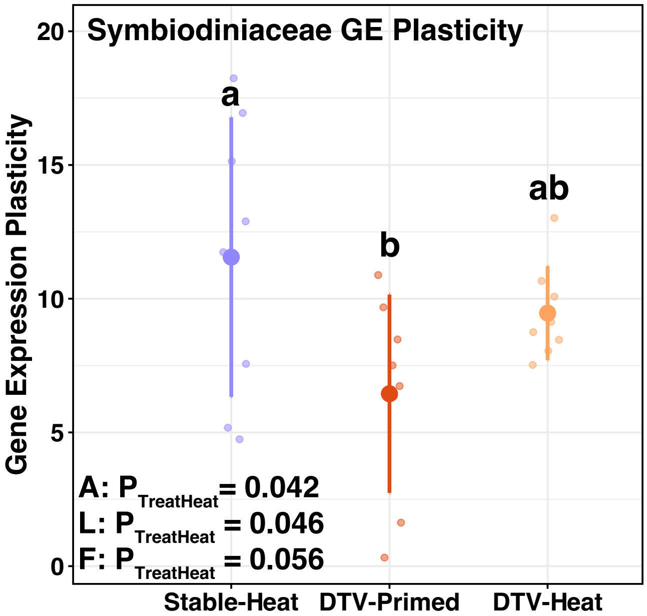


**Figure S8. Symbiodiniaceae gene expression (GE) plasticity.** Mean GE plasticity and standard deviation by experimental treatment using the "Stable-Primed" group as a control. Significance was determined by two-way ANOVA and Tukey HSD test (lowercase letters) for differences across treatments. For differences in variance Levene's Test (L) and Fligner's Test (F) were used with significant pairwise comparisons detailed in the text.

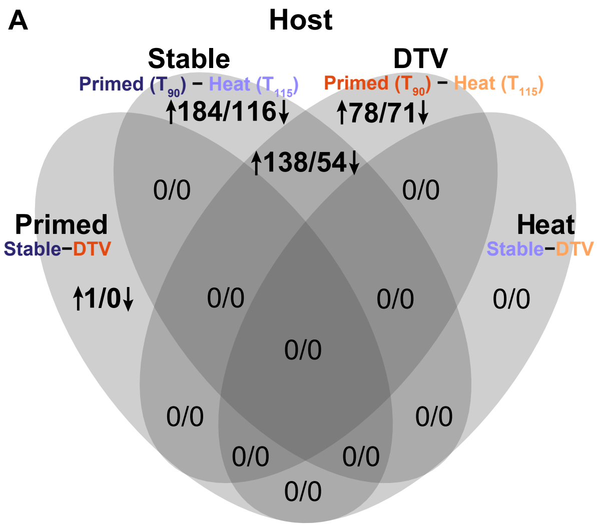


**Figure S9. Overlap of differentially expressed genes (DEGs, adjusted p-values < 0.05) across thermal priming and heat challenge.** A) Combined host gene expression and B) Symbiodiniaceae gene expression by treatment groupings. Upregulated genes (↑: Padj < 0.1, log2 fold change > 0) are more highly expressed in the first group listed in the contrast relative to the second. Conversely, downregulated genes (↓: Padj < 0.1, log2 fold change < 0) show lower expression in the first group listed in the contrast relative to the second.

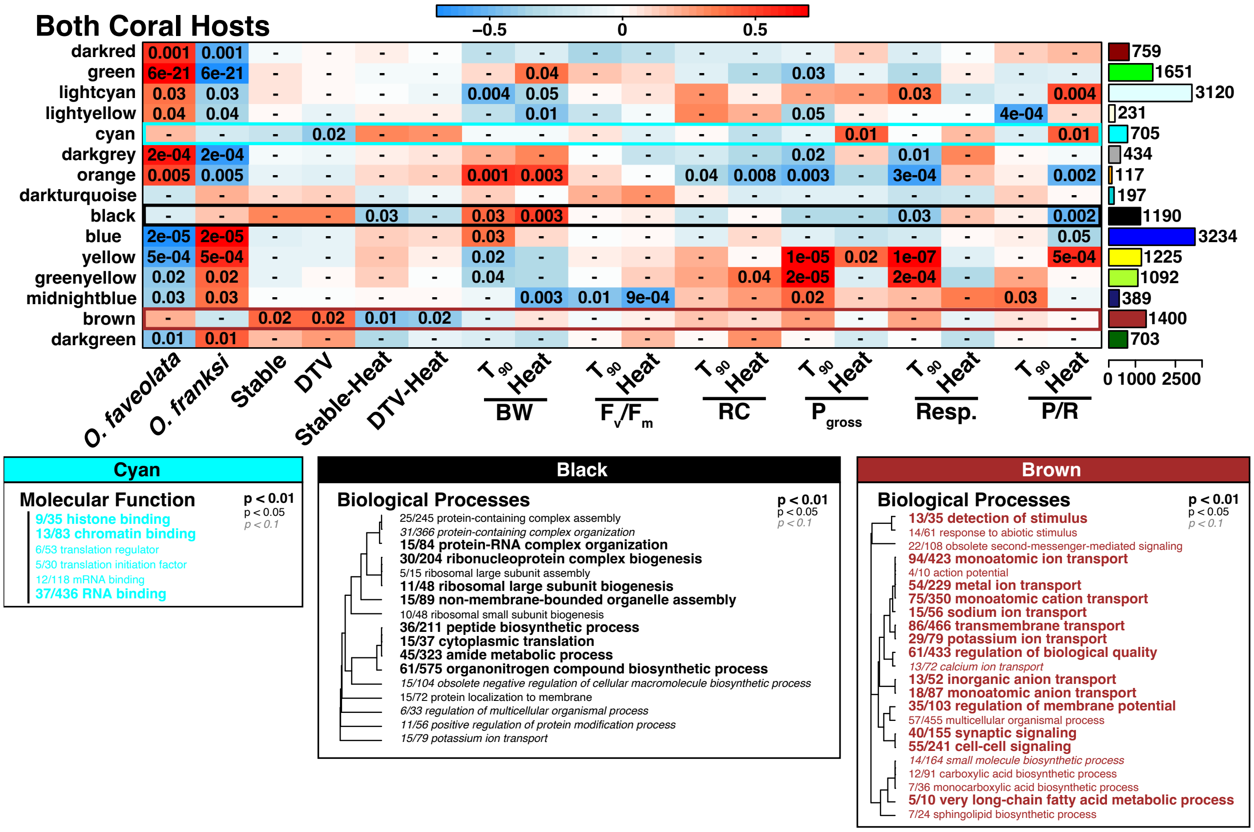


**Figure S10. Weighted gene co-expression network analysis (WGCNA) for combined host analysis across thermal priming and heat challenge.** Module eigengene trait relationships identified by WGCNA with correlation values ranging from 1 (red) to -1 (blue) and associated significant p-values are included within heatmap cells. Modules significantly correlated with treatments (Stable-Primed, DTV-Primed, Stable-Heat, DTV-Heat) are indicated by boxes in the module color and GO terms enriched within these modules shown below the module-trait heatmap (cyan, black, brown) as dendrograms. Enriched Gene Ontology (GO) terms in the molecular function (MF) and biological process (BP) categories are shown. Font size and italics indicate levels of statistical significance as indicated by the inset key. Dendrograms show sharing of genes between GO terms and the fractions indicate the proportion of genes contained within the module relative to the total number of genes within the GO category.

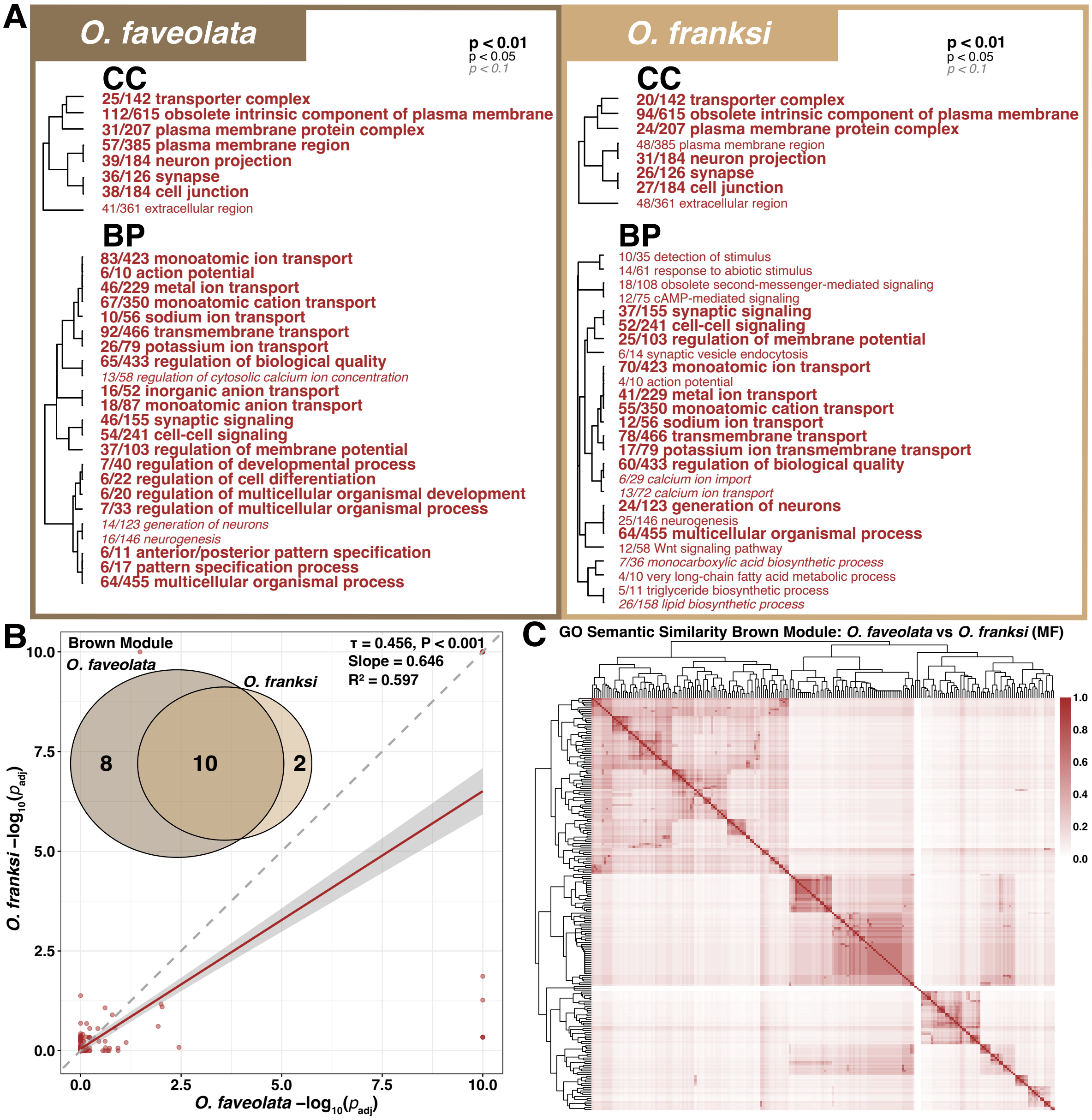


**Figure S11. Comparisons of Brown Modules identified by WGCNA across the two *Orbicella* host species.** A) Dendrograms of Gene Ontology (GO) terms enriched in the brown modules identified by the WGCNA for *O. faveolata* (left) and *O. franksi* (right) hosts. Enriched terms in the cellular component (CC) and biological process (BP) categories are shown. Font size and italics indicate levels of statistical significance as indicated by the inset key. Dendrograms show sharing of genes between GO terms and the fractions indicate the proportion of genes contained within the module relative to the total number of genes within the GO category. B) Comparison of -log10(Padj) values for each enriched GO term shared between the brown modules WGCNA for *O. franksi* and *O. faveolata* with a Kendall correlation (τ) determining the significance and correlation of enrichment. Insert is the overlap of significant GO terms identified from each brown module between the two host species. B) GO term heatmap showing semantic similarity of GO terms within each host's brown module. Rows and columns are GO terms from both brown modules, with terms grouped by similar biological meaning of molecular function GO terms. Brown color indicates higher Wang semantic similarity and white indicates no similarity.

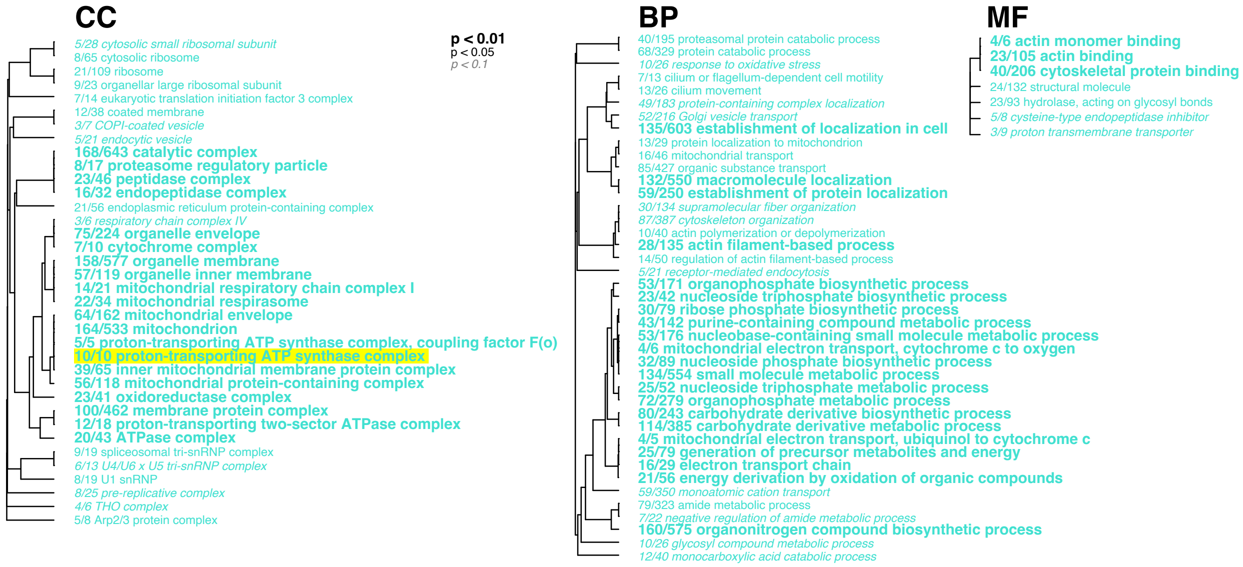


**Figure S12. Hierarchical clustering of Gene Ontology (GO) terms enriched in the paleturquoise module of the *O. franksi* host WGCNA analysis.** Dendrograms of GO terms enriched in the paleturquoise module identified by the WGCNA for *O. franksi* hosts. Enriched terms in the molecular function (MF), cellular component (CC) and biological process (BP) categories are shown. Font size and italics indicate levels of statistical significance as indicated by the inset key. Dendrograms show sharing of genes between GO terms and the fractions indicate the proportion of genes contained within the module relative to the total number of genes within the GO category. A parent term specifically associated with V- and F-type ATPases is highlighted.
